# Functionalizing cell-free systems with CRISPR-associated proteins: Application to RNA-based circuit engineering

**DOI:** 10.1101/2021.04.08.438922

**Authors:** François-Xavier Lehr, Alina Kuzembayeva, Megan E. Bailey, Werner Kleindienst, Johannes Kabisch, Heinz Koeppl

## Abstract

Cell-free systems have become a compelling choice for the prototyping of synthetic circuits. Many robust protocols for preparing cell-free systems are now available along with toolboxes designed for a variety of applications. Thus far the production of cell-free extracts has often been decoupled from the production of functionalized proteins. Here, we leveraged the most recently published protocol for *E. coli*-based cell extracts with the endogenous production of two CRISPR-associated proteins, Csy4 and dCas9. We found pre-expression did not affect the resulting extract performance, and the final concentrations of the endonucleases matched the level required for synthetic circuit prototyping. We demonstrated the benefits and versatility of dCas9 and Csy4 through the use of RNA circuitry based on a combination of single guide RNAs, small transcriptional activator RNAs and toehold switches. For instance, we show that Csy4 processing increased fourfold the dynamic range of a previously published AND-logic gate. Additionally, blending the CRISPR-enhanced extracts enabled us to reduce leakage in a multiple inputs gate, and to extend the type of Boolean functions available for RNA-based circuits, such as NAND-logic. Finally, the use of dual transcriptional and translational reporters for the engineering of RNA-based circuits, allowed us to gain better insight into their underlying mechanisms. We hope this work will facilitate the adoption of advanced processing tools for RNA-based circuit prototyping in a cell-free environment.

## Introduction

Cell-free expression systems have become an established tool for bottom-up approaches in synthetic biology. By decoupling the engineering of bio-molecular modules and systems from the whole cell complexity, cell-free gene expression (CFE) technology has proven useful for a variety of purposes from understanding basic cellular processes to biomanufacturing.^1^ Beyond bulk protein production, recombinant and crude extract systems have enabled rapid developments in biosensing, model-driven design of gene circuits, high-throughput library screening, and prototyping of circuits based on a large repertoire of characterised parts.^1,2^

CFE system based on crude extracts can be made from a variety of cell types, but *Escherichia coli* (*E. coli*) is the most popular because of its ease of production and high yield at a lower overall cost. ^3^ Another advantage of the crude cell extracts is that the native *E. coli* machinery can be used for transcription with a high-yield. ^4^ The protocol for producing CFE extract has been streamlined and can now be performed in 1-2 days. Energy systems are also continuously revised to account for new information from metabolic pathways on optimization of ATP regeneration and improving the overall productivity and longevity of the reactions. ^5^

Recently, Silverman et al. ^6^ has deconstructed the effects of diverse protocols in order to standardize a robust, engineerable and low-cost CFE extract preparation. Synthetic biology is now harnessing the power of CFE extracts for a variety of purposes. In particular, circuit prototyping has taken advantage of the CFE extract ability to process large libraries of components quickly.^7^ The continuous increase in both circuit complexity and functionality translates into a higher metabolic burden for the CFE extract. Circuit behavior can be impacted by transcription and translation competition if concentrations of parts are close to saturation. ^8,9^ For instance, it has been shown that co-expressing proteins in CFE system creates more burden to the translational machinery and impact the final concentration of the translated components negatively. ^10^

One solution to alleviate the CFE extract burden is to use purified components to functionalize the environment. This has been used for instance with the supplementation of proteins enabling the use of linear DNA,^11^ or to adjust the DNA or RNA degradation rates. ^12^ However, the requirement of multiple functions involves the use of multiple exogenous proteins. High concentrations of buffering compounds or additives present in the protein storage buffer can be detrimental to the CFE yield. Using protein from multiple batches as well as increased liquid-handling operations could also contribute to diminishing the reproducibility across experiments.^13^ Finally, the purification of additional proteins may present additional challenges, such as low protein recovery yields, and can ultimately lead to delayed access to CFE prototyping. ^14^ One solution to this problem is to pre-express the protein of interest in the bacterial strain prior to producing the extract. Recently, Soye et al. ^15^ expressed T7 RNA polymerase in a genomically recoded strain and demonstrated high yield in the resulting CFE extract. Dudley et al. ^16^ mixed multiple lysates enriched with different overexpressed enzymes to reconstruct the full mevalonate pathway. This highly flexible approach was also applied to n-butanol biosynthesis^17^ and site-specific glycosylation of target proteins. ^18^ However, this one-pot strategy has been principally restricted to metabolic engineering and bioproduction purposes, but still needs to be extended to circuit prototyping in synthetic biology.

In this work, we combine the standardized protocol from Silverman et al. ^6^ and the expression of two CRISPR-associated proteins in the BL21 strain to produce CFE extracts with augmented functions. Marshall et al. ^19^ and Liao et al. ^20^ have recently shown the great capacity of CFE platforms for characterizing a variety of CRISPR-Cas systems. Here, we first show that the augmented CFE extracts retain a high-yield efficiency compared to the standard CFE extract. We then characterise the two CRISPR-systems: Cas9 endonuclease dead (dCas9) with the engineering and testing of different small guide RNAs as well as Csy4 endoribonuclease with the cleavage of either Ribosome Binding Sequences (RBS) or 5’ Untranslated Regions (UTRs) from the gene mRNA.

We demonstrate how circuit prototyping, in particular when applied to RNA-based devices, can benefit from the multiplexed processing of dCas9 and Csy4. We use our one-pot strategy to specifically engineer or enhance modules based on RNA-regulators, featuring orthogonality, low metabolic burden, and fast signal propagation. Small Transcriptional Activator RNAs (STARs) and toehold switches characterised from our previous work ^21^ were combined with single-guide RNAs (sgRNA) and Csy4 insulating hairpins to improve Boolean logic processing. We also gained insight into the transcriptional state of the AND-gate using Malachite Green Aptamers (MGAs), which may also reduce the uncertainty of inferred parameters when predictive computational models are required. ^10,22^ Overall, we hope this work will facilitate the adoption of augmented CFE extracts where a broad range of applications, from molecular diagnostics to educational toolkit, ^23^ could take advantage of the embedded CRISPR features. Finally, we anticipate this one-pot CFE strategy could facilitate and accelerate the access to advanced circuit processing tools for small-sized laboratories by skipping additional protein purification steps.

## Results and discussion

### Pre-expressing proteins does not affect cell-extract performance

The CFE extracts were prepared mainly according to the protocol described by Silverman et al. ^6^ for BL21 Rosetta 2 strain. We made few adaptations: both the dialysis and the flash freezing of the cell pellet were omitted, to constrain the preparation time to one day. Our data suggest the dialysis step did not improve the CFE yield (Supp. Fig. 1), in agreement with previous published CFE protocols tested with bacteriophage-based promoters. ^10,24^ However, we note that the protocol from Silverman et al. ^6^, tested with a native promoter, reported a higher yield in the dialysed CFE extract (~ 30% more) compared to the run-off alone. This result may encourage the operator to explore this parameter considering the large variability reported between extracts prepared from various laboratories. ^25^

First of all, we aimed to compare the efficiency of the control CFE extract (denoted as wild-type CFE extract from here forward) with three different CFE extracts pre-expressing the proteins of interest (denoted as augmented CFE extracts from here forward). Briefly, an ampicillin resistance plasmid expressing mRFP1, dCas9, or Csy4 downstream of the medium-strength *E. coli* promoter J23108 were transformed into BL21 Rosetta 2 strain and prepared according to the standard protocol (Fig. 1). Our data suggest that the doubling time of the enhanced strain was not affected by the supplementary plasmid. The augmented and WT CFE extracts were all able to reach *OD*_600_ = 3 after 4 to 5 hours of culture. (Supp. Fig. 2). We measured the overall protein concentration via Bradford assay in WT lysates at different ODs and in two different enhanced lysates (Supp. Fig. 3. Our data indicate all resulting protein concentrations were contained in the typical range (20-40 mg/ml) obtained from similar CFE systems ^26,27^. The WT and enhanced lysates were treated similarly and supplemented with the same energy regeneration buffer. A reporter plasmid, Pr-sfGFP-MGA4x, was used in the following experiments to assess the CFE extract efficiency. The superfolder green fluorescent protein (sfGFP) was used to monitor translation, and four tandem repeats of Malachite Green Aptamers (MGA) enabled the monitoring of transcription. ^28^ Examples of fluorescence time-courses for 1 nM of reporter plasmid are displayed in Fig. 1 A. The sfGFP levels reached a stable plateau after several hours of expression while the mRNA signal reached a transient maximum after a couple of hours. The subsequent MGA decreasing signal reflects the mRNA degradation in the CFE extract. ^10^ Titration of the reporter plasmid showed similar transcription and translation dose-response in the four CFE extracts (Fig. 1 B). Both processes reached their maximum yield at around 20 nM of the input reporter plasmid. Purified sfGFP-MGA4x mRNA and eGFP were used for calibrating the transcriptional and translational yield, respectively (Supp. Fig. 4). The yields (up to 0.2-1 μM of mRNA and 1-5 μM of sfGFP) are comparable to those achieved by other groups. ^6,24^ This suggests that functionalizing CFE extracts with pre-expressed protein does not have a noticeable impact on the final output. To confirm this trend, we performed statistical analysis to compare the yields of WT CFE extracts with the yields of enhanced CFE extracts. The yields in mRNA and sfGFP production were not found to be significantly different between three independently produced WT CFE extracts and six enhanced CFE extracts (Supp. Fig. 5).

**Figure 1:**
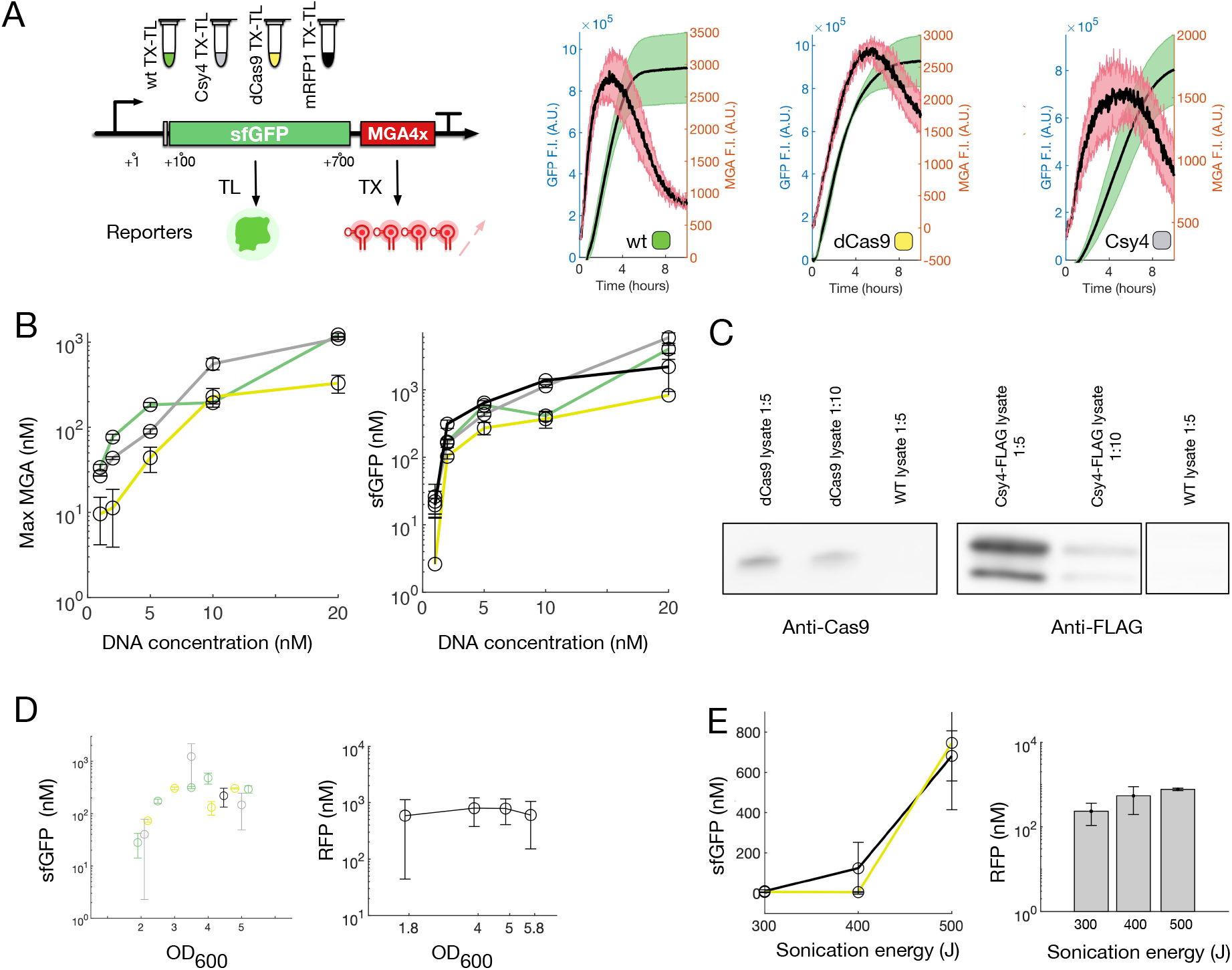
Influence of the pre-expression of proteins in BL21 rosetta™ 2 cells on the resulting CFE efficiency. A) The standard reporter plasmid pr-sfGFP-MGA4x enables simultaneous monitoring of the transcription and translation production. Four different CFE extracts are considered here: the wild-type (WT), and three CFEs pre-expressing Csy4 (grey), dCas9 (yellow), or mRFP1 (black). Examples of sfGFP (green) and MGA (red) fluorescence traces for the WT, Cys4 and dCas9 CFE extracts expressing 1 nM of reporter plasmid.B) sfGFP (right) and MGA (left) productions in WT and enhanced CFE extracts titrated with the reporter plasmid. Error bars represent the standard deviation of the mean from three technical replicates. C) Western blot analysis of dCas9 and Csy4-FLAG activators demonstrating that constructs are effectively translated in the lysates. Ratios indicate crude extract:buffer dilution factors. D) (left) sfGFP yields of the CFE extracts for different *ODs*_600_ with 1 nM of reporter plasmid. (right) pre-expressed RFP concentrations in mRFP1 CFE extracts for different *ODs*_600_. E) Influence of the sonication energy on the CFE performance for mRFP1 and dCas9 CFE extracts (1 nM of reporter plasmid).

We also demonstrated the presence of the pre-expressed proteins in the CRISPR-enhanced extracts via western blot analysis (Fig. 1 C, Supp. Fig. 6, Supp. Fig. 8). We estimated the final concentration of dCas9 in the lysate to be ~ 68±32 nM (Supp. Fig. 7). For Csy4, a rough quantification via Coomassie Blue staining gave us a final concentration in the range of 4.4 μM - 8.8 μM (Supp. Fig. 8). Additionally, we obtained an indirect quantification of the pre-expressed protein in an enhanced extract expressing mRFP1 by using a calibration curve (Supp. Fig. 4). mRFP1 lysates harvested from mid to late exponential phase (*OD*_600_ = 2-6) displayed a similar quantity of mRFP1, up to ~ 0.8 μM (Fig. 1 D right). The difference of magnitude in the final concentration of Csy4 and dCas9 could be explained by their difference in coding size and/or complexity of transcription/translation. For the corresponding *OD*_600_, the CFE extracts showed a typical sfGFP dose-response yield, with a maximum reached at around *OD*_600_ = 3, as reported previously by Kwon and Jewett 24 (Fig. 1 D left). Finally, we tested whether the sonication energy led to a variation in the final amount of pre-expressed protein (Fig. 1 E right). We found no difference in the amount of pre-expressed mRFP1 between the subjected sonication energies (300 - 500 J). Conversely, the sfGFP productivity of the CFE extract was highly dependent on the sonication energy, in agreement with previous reports^10,24^ (Fig. 1 E left).

### Csy4 endoribonuclease as a tool to process mRNA in TXTL

Csy4 is a bacterial endoribonuclease that comes from *Pseudomonas aeruginosa* that can be utilized for RNA processing. ^29^ It recognizes and cleaves an RNA substrate at a specific 28-nt hairpin sequence, cleaving at the 3’ end of the hairpin. We chose to create a CFE extract with Csy4 pre-expressed in order to utilize the RNA processing capabilities of the endoribonuclease. The J23108 promoter (relative strength = 0.5 according to the Anderson promoter collection chosen as a reference standard ^30^) was utilized to limit the over-expression of the protein in the cell extracts, which may lead to toxicity effects towards the host cell or to the creation of inclusion bodies detrimental for downstream applications. ^31^

#### Csy4 controls translation in a tandem of fluorescent proteins

To assess the cleavage efficiency of the pre-expressed Csy4 protein, we used a tandem protein fusion composed of sfGFP and mCherry driven by a strong J23119 *E. coli* promoter. Two constructs were cloned, differing by a Csy4 hairpin inserted downstream to the RBS of mCherry (Fig. 2 A). Upon cleavage of the hairpin by the endoribonuclease, mCherry mRNA translation should be inhibited due to the absence of upstream RBS. As a control experiment and to compare with previous similar *in vivo* experiments, we first tested the system in *E. coli*. Briefly, cells were co-transformed with or without plasmid expressing Csy4 and with either the sfGFP-Csy4-mCherry or sfGFP-mCherry plasmids. In the absence of co-expressed Csy4, the mCherry signal from sfGFP-mCherry was found to be about 2-fold higher than the one measured for sfGFP-Csy4-mCherry (Fig. 2 B). This difference could be explained by the additional translated amino acids of the Csy4 hairpin sequence. To avoid any bias, the lower level of Csy4-mCherry fluorescence will be taken into account to quantify the effect of the Csy4 endoribonuclease alone. In the presence of Csy4 protein, mCherry intensity difference increased to 12.5-fold between the two constructs. The sharp decrease of the sfGFP-Csy4-mCherry fluorescence was close to the baseline level, which supports the data of RBS cleavage experiments published by Qi et al. ^29^. Interestingly, sfGFP-Csy4-mCherry showed 4-fold higher sfGFP intensity than sfGFP-mCherry in the absence of Csy4 (Fig. 2 B). Upon Csy4 expression, sfGFP-Csy4-mCherry displayed a sharp decrease in sfGFP expression compared to the sfGFP-mCherry construct (2-fold lower). We hypothesized the Csy4 hairpin, inserted downstream to the sfGFP sequence, could either increase the mRNA stability or act as a high-performing insulator. The resulting data can also be visualized on heatmaps, where the intensity values of sfGFP-Csy4-mCherry were normalized to sfGFP-mCherry in each extract condition (Fig. 2 C). In other words, a value close to 0 suggests a complete inhibition of the fluorescent protein, while a value of 1 indicates no difference between the two constructs.

**Figure 2:**
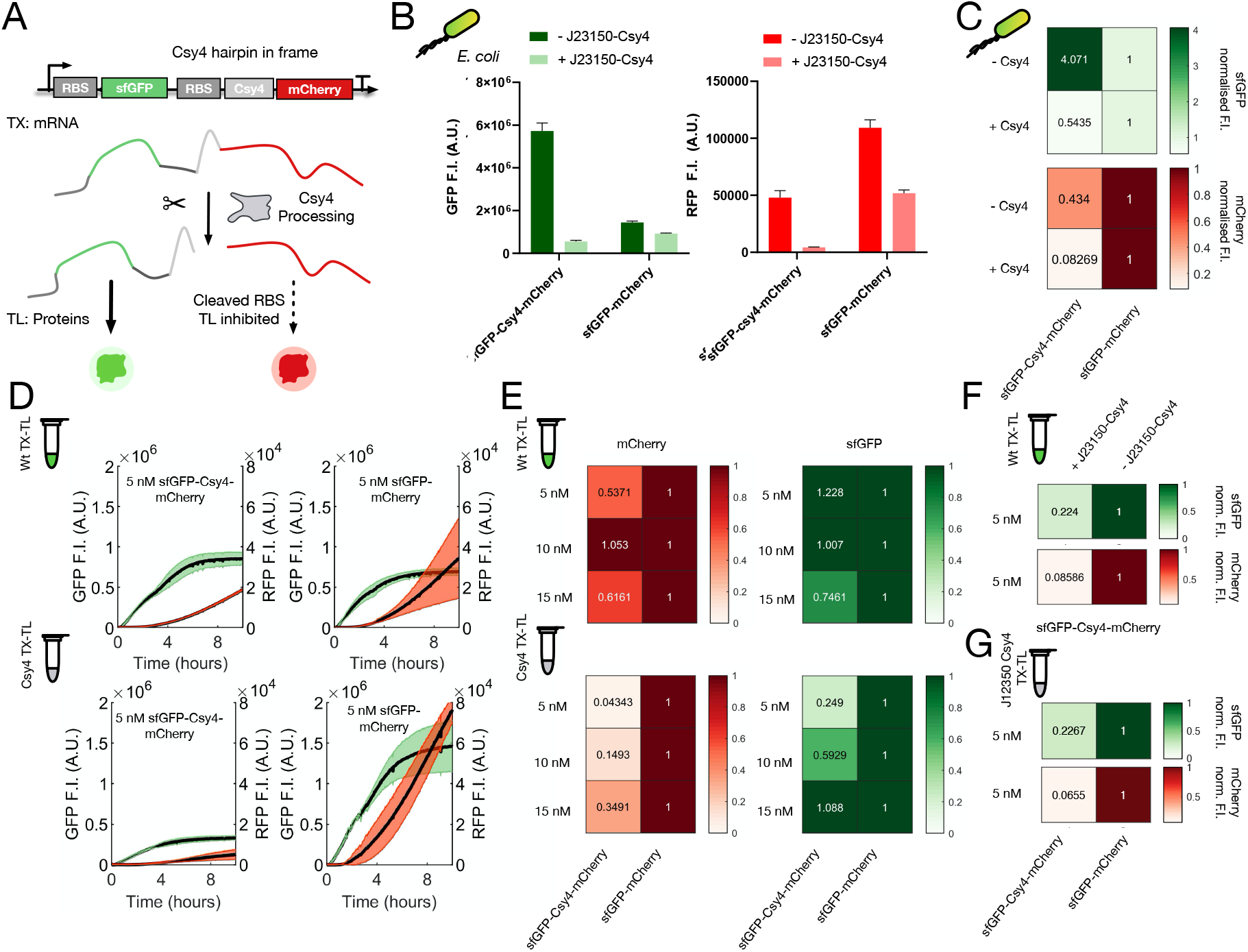
Csy4 controls gene expression in tandem of fluorescent proteins. A) Scheme of the Csy4 processing mechanism applied to an operon composed of sfGFP and mCherry. Upon cleavage of the hairpin by Csy4, mCherry mRNA is separated from the RBS. B) Implementation *in vivo*. Fluorescence measurements of sfGFP-Csy4-mCherry and sfGFP-mCherry in the presence or the absence of Csy4 encoding plasmid. C) Normalized mCherry and sfGFP fluorescence heatmaps computed from *in vivo* measurements. D) mCherry (left axis) and sfGFP (right axis) time-courses in Csy4 and WT extracts corresponding to 5 nM of either sfGFP-Csy4-mCherry or sfGFP-mCherry. E) Normalized mCherry and sfGFP fluorescence heatmaps computed from *in vitro* reactions at three different DNA concentrations. F) Normalized mCherry and sfGFP fluorescence heatmaps for 5 nM of sfGFP-Csy4-mCherry in WT extract supplemented with 5 nM of Csy4 encoding plasmid. G) Normalized mCherry and sfGFP fluorescence heatmaps in a J23150-Csy4 extract for 5 nM of either sfGFP-Csy4-mCherry or sfGFP-mCherry. Error bars and shaded regions represent the standard deviation of the mean from three technical replicates.

In the next step, we transferred the sfGFP-mCherry fusion experiment to CFE systems. We used two CFE extracts: a WT extract (with no Csy4 protein) and a Csy4 extract (pre-expressing Csy4 from a J23108 promoter). Csy4 lacks the ability to engage in multiple-turnover catalysis. ^32^ Consequently, we expected to observe the saturation of the cleavage activity at increasing concentration of mRNA, which can be indirectly controlled with DNA concentration (Fig. 1 D). To test this hypothesis, we titrated the two tandem constructs at three concentrations (5 nM, 10 nM, 20 nM) in both extracts. Kinetic traces are reported in Fig. 2 D and Supp. Fig. 9. The corresponding heatmaps are displayed in Fig. 2 E and used the same normalization as *in vivo*. In the WT extract, sfGFP-Csy4-mCherry displayed normalized mCherry fluorescence values between 0.53 and 1. These values are in the same order of magnitude as the one found in vivo (0.43). That suggests the detrimental impact of the additional Csy4 amino acids observed *in vivo* transfers similarly *in vitro*. In the Csy4 extract, sfGFP-Csy4-mCherry displayed normalized mCherry values as low as 0.04 (for 5 nM of plasmid) and 0.15 (for 10 nM of plasmid). Those values match again the data found *in vivo* in cleavage condition (0.08). At 20 nM of plasmid, the normalized mCherry value increased to 0.34 and is thus closer to the values computed in non cleavage condition both *in vivo* and *in vitro*. Taken together, these data suggest efficient cleavage activity below 10 nM of DNA template and incomplete cleavage of the mRNA transcripts at 20 nM of template. The sfGFP normalized heatmaps corroborate this observation. In the WT extract, we found sfGFP normalized values to be in the same order of magnitude between the two constructs (0.75 - 1.23). In the Csy4 extract, sfGFP normalized ratios increased with plasmid concentration (0.25, 0.59, and 1 for 5, 10, and 20 nM of plasmid, respectively). The inhibition observed at 5 nM and 10 nM is weaker than mCherry, but this reflects the data shown *in vivo* and supports the hypothesis suggesting that Csy4 is saturated above 10 nM of DNA template. Additionally, we produced another Csy4 CFE extract with a weaker promoter (relative strength = 0.2). In this extract, we observed a similar sfGFP and mCherry inhibition as in the J23108-Csy4 CFE extract for 5 nM of the fusion constructs (Fig. 2 G, Supp. Fig. 10 B). Finally, we performed an additional experiment to test whether Csy4 can be co-expressed during the cell-free reaction and to compare the cleavage performance with our augmented extract. In a WT extract, we co-expressed 5 nM of sfGFP-Csy4-mCherry with 5 nM of a plasmid expressing either Csy4 or a decoy protein from a J23150 constitutive promoter (Fig. 2 F, Supp. Fig. 10 A). The resulting data were similar to the in vitro and *in vivo* characterization established previously for 5 nM of plasmid and showed strong inhibition of mCherry (normalized value of 0.08) and sfGFP (normalized value of 0.15).

#### Cleavage of 5’ UTRs inhibits gene expression in cell-free systems

We performed a second investigation of Csy4 cleavage activity in the augmented extract, to test whether different experimental conditions may affect the optimal DNA range previously determined. Two strong 5’ UTRs (ybaB-AG, lrp) were chosen among different 5’ UTR constructs based on the Pr-sfGFP-MGA4x reporter plasmid (Supp. Fig. 11). Two additional constructs were made by inserting a Csy4 hairpin between the 5’ UTR and the sfGFP start codon (ybaB-AG-Csy4, lrp-Csy4). In the presence of Csy4 endoribonuclease, the Csy4 hairpin should be cleaved and the 5’ UTR separated from the sfGFP mRNA (Fig. 3 A). Consequently, we expect the mRNA translation rate to become independent from the 5’ UTR. Previous experiments in living cells have shown that various 5’ UTR constructs exhibit homogeneous production rates after cleavage. ^29^ However, it is not yet known how well 5’ UTR cleavage transfers into CFE systems. A fifth construct had its 5’ UTR replaced with the downstream sequence to the cleavage site of Csy4 hairpin (5’-CTAAGAA - 3’). This construct will be valuable to measure the expected behavior of the cleaved UTRs in the CFE extracts. We titrated all constructs in the Csy4 and WT extracts from 1 nM to 20 nM. Examples of kinetic traces can be found in Supp. Fig. 12 B. We normalized the endpoint fluorescence data to our standard reporter construct expressed in each extract (Supp. Fig. 12 A). The corresponding sfGFP heatmaps are displayed in Fig. 3 B. The construct mimicking a cleaved 5’ UTR expressed poorly at concentrations below 10 nM in the WT and Csy4 extracts. Interestingly, Silverman et al. ^6^ found no expression from a construct lacking a stability hairpin in its 5’ UTR, which could explain our control construct’s poor performance. This is reinforced by the low level of MGA fluorescence achieved by the cleaved mimic (Supp. Fig. 12 B). In the WT extract, all constructs with and without Csy4 hairpin expressed a typical^33^ increase of gene expression with DNA concentration. The ybaB-AG and lrp constructs displayed comparable sfGFP dose-responses in the Csy4 extract. Strikingly, in the Csy4 extract, ybab-AG-Csy4 and lrp-Csy4 displayed low normalized values for 1 to 10 nM of DNA, and showed a subsequent increase of gene expression at 20 nM. Their dose-responses are thus similar to those obtained for the construct mimicking the 5’ UTR cleavage, suggesting cleavage of the 5’ UTR up to 10 nM of the DNA template. The data suggest however a higher cleavage performance at 20 nM of DNA template compared to the fusion protein experiment. We are not able to directly explain this observation, but we hypothesize that the Csy4 binding hairpin could be heavily constrained by its flanking sequences, as shown by Guo et al. ^34^.

**Figure 3:**
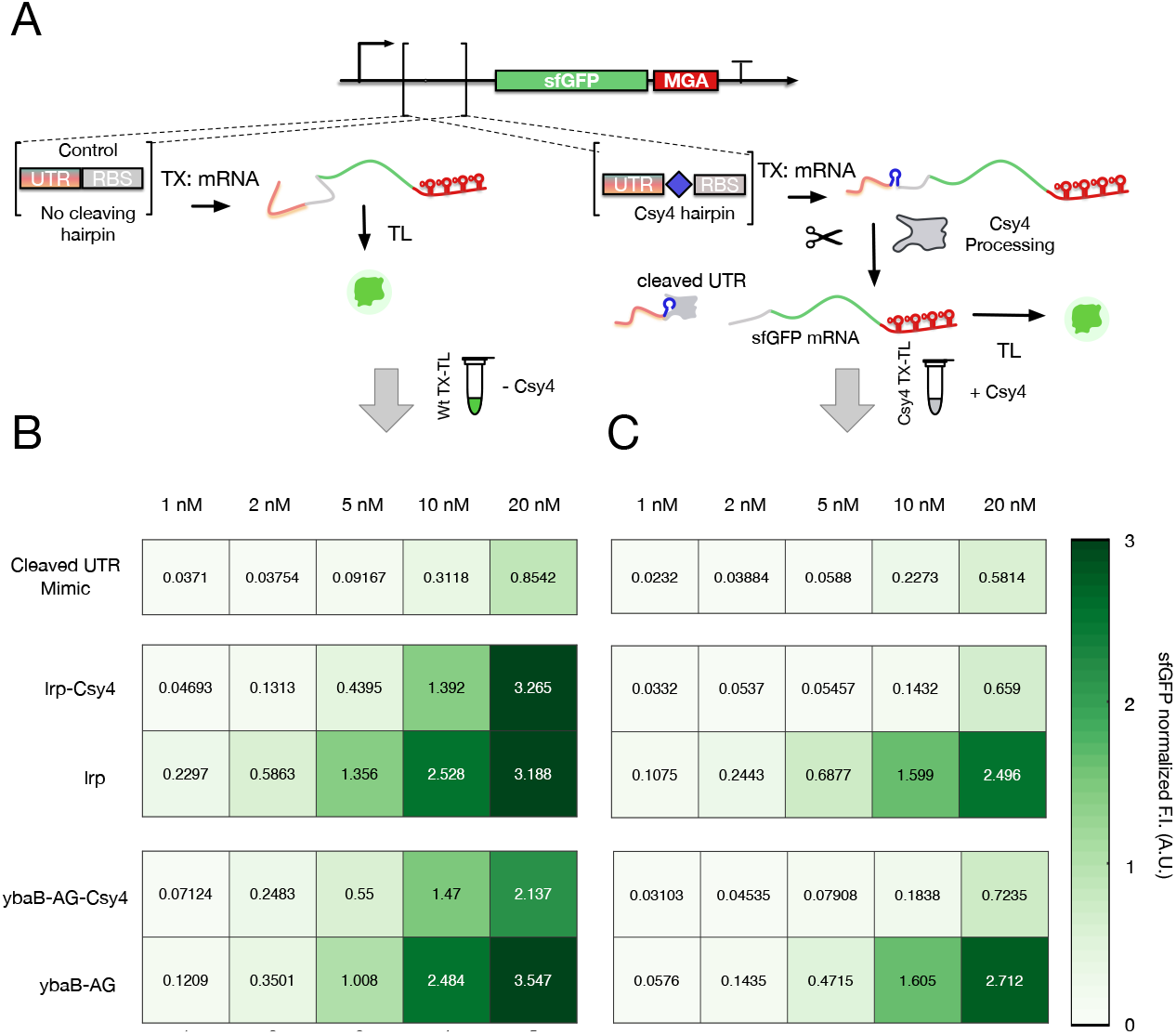
5’ UTR cleavage decreases gene expression in augmented Csy4 CFE extract. A) Scheme of the Csy4 processing mechanism. SfGFP reporter constructs contain either 5’ UTRS without Csy4 hairpin, or are separated from the sfGFP coding sequence and the RBS by a Csy4 hairpin. Upon cleavage of the hairpin by Csy4 endoribonuclease, the UTR is separated from the mRNA. B) Normalized sfGFP heatmaps of five UTR constructs titrated between 1 and 20 nM in WT CFE extract. C) Normalized sfGFP heatmaps of five UTR constructs titrated between 1 and 20 nM in Csy4 CFE extract. Error bars and shaded regions represent the standard deviation of the mean from three technical replicates.

Csy4 has been shown to remain bound to the processed mRNA,^32^ so we also asked if the Csy4-mRNA complex affects mRNA stability, as it could be beneficial for RNA-circuits. We measured the decay rate of purified MGA4x mRNA in the Csy4 CFE extract with and without Csy4 hairpins placed at the 5’ and 3’ ends, but the data did not suggest difference in their half-lives (Supp. Fig. 14). Taken together, the data demonstrate functional Csy4 cleavage in our augmented Csy4 extract. The cleavage activity seems optimal for a DNA concentration compatible with many prototyping applications in TXTL, as most of the parts are often set between 0.5 nM to 10 nM. ^12,19,35^

### Characterisation of dCas9 activity in CFE extract

To assess the pre-expressed dCas9 activity in the corresponding augmented CFE extract, four single guide RNAs (sgRNAs) were designed to maximize the ON-target score and minimize the OFF-target score. ^36^ Three sgRNAs bind in the non-target strand of the sfGFP coding sequence (5’, middle and 3’ of the gene, denoted as sg-1,-2,-3) while the fourth sgRNA binds in the J23119 promoter sequence (sg-4). In the presence of the sgRNAs, the sfGFP production should be inhibited since the transcription elongation is hindered by the dCas9-sgRNA complex blocking the DNA template (Fig. 4 A). Both sfGFP and MGA fluorescence signals are therefore expected to be lowered in the presence of the sgRNA.

**Figure 4:**
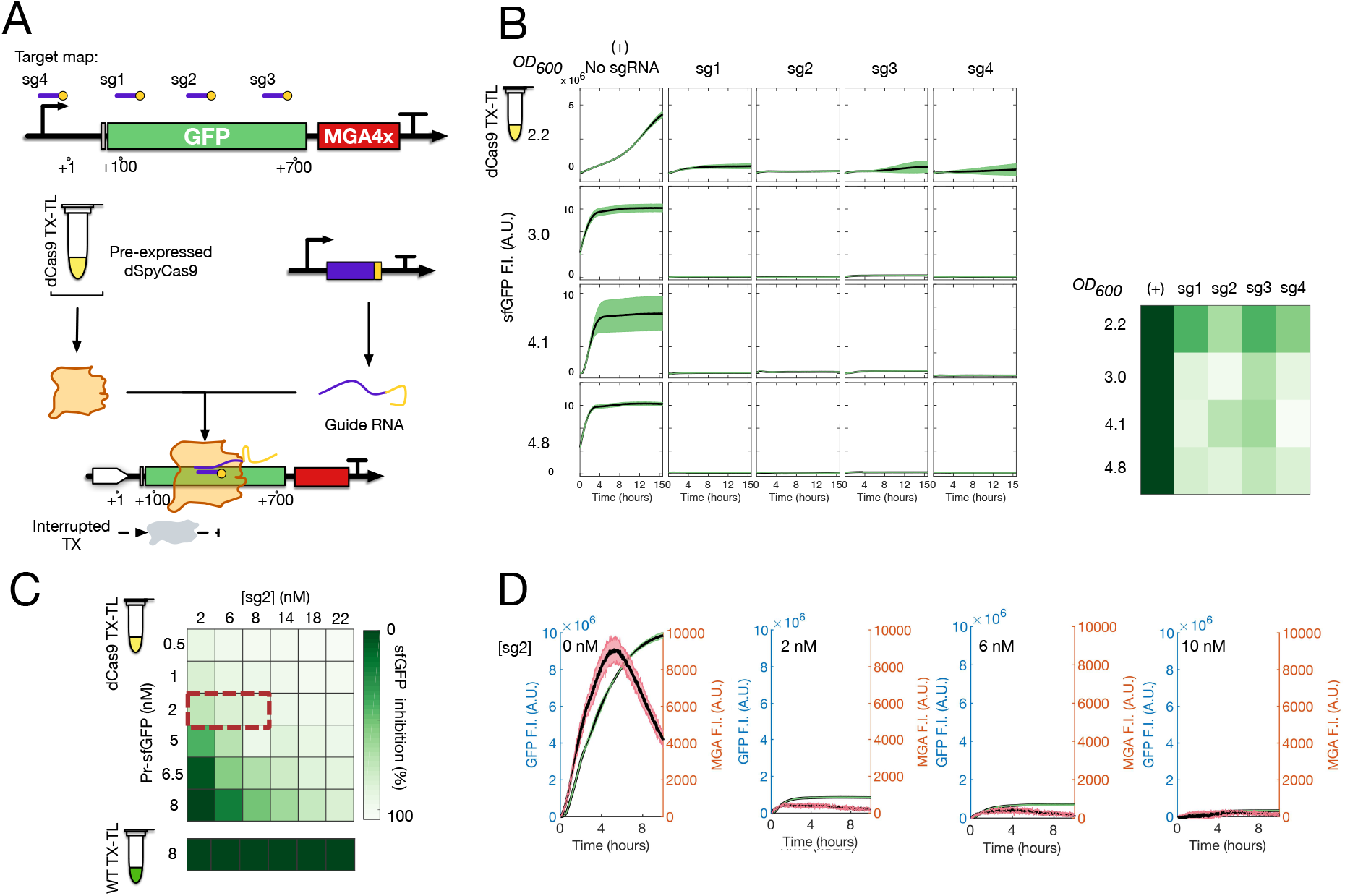
dCas9 regulates gene expression in enhanced CFE extracts. A) Scheme of the dCas9 regulation mechanism and the template target location of the four different sgRNAs. B) sfGFP time traces for different sgRNAs in CFE extracts made from various *ODs*_600_ (left). Corresponding heatmap summary of the sfGFP production rates (right). C) Dose-response matrix of sfGFP production rates from reactions titrated with both reporter template and sgRNA plasmid. D) Sample of sfGFP (green) and MGA (red) fluorescence time traces (corresponding to the sgRNA concentrations framed by the red dashed line in D)). Error bars represent the standard deviation of the mean from three technical replicates.

#### dCas9 is active regardless of the CFE extract harvest time

Four dCas9 CFEs made from different harvest time spanning from a range of mid to late exponential growth (*OD*_600_ = 2.2, 3.0, 4.1 and 4.8) were used to test the activity of dCas9 along with 10 nM of each sgRNA plasmid and 1 nM of the reporter plasmid. Fluorescence time traces and production rates are shown in Fig. 4 B and Supp. Fig. 15. In the WT CFE extract, we observed similar kinetic traces for MGA and sfGFP fluorescences between the control experiment and the four different sgRNAs, showing no inhibition in the absence of dCas9 (Fig. supp. 16).

In the dCas9 CFE extracts, sfGFP kinetics show a dramatic decrease in fluorescence for all conditions, regardless of the *OD*_600_ or the sgRNA. This suggests that the dCas9 concentration is similar among extracts processed at different *OD*_600_. However, the dynamic range is one order of magnitude lower in the CFE extract made from *OD*_600_ = 2.2, which is likely due to the lower sfGFP productivity of the cell extract made from cells harvested at the early exponential phase (Fig. 1 B). The sgRNAs sg-1, sg-2 and sg-4 display a similar sfGFP inhibition, while sg-3 consistently exhibit a reduced dynamic range. This result may be explained by the DNA target of sg-3, located towards the 3’ end of the sfGFP coding sequence, enabling the transcription of a near-to-complete sfGFP mRNA.^19^ The MGA production rate does not show any obvious correlations between sgRNA targets or *OD*_600_, likely due to the low signal-to-noise ratio (SNR) of the transcribed mRNA. To fully characterize the sgRNA behavior, we titrated the reporter template and sg-2 with plasmid concentrations relevant to cell-free prototyping^12^ (Fig. 4 C). While no inhibition was observed in the WT CFE extract for the range of sg-2 concentrations at 8 nM of reporter plasmid (see Supp. Fig. 17 for fluorescent time traces), a correlation between the two DNA components was observed in the dCas9 CFE extract. Under 2 nM of template, the plasmid was repressed with even low concentrations of sgRNA plasmid. Above 2 nM, an increasing concentration of sgRNA plasmid was required to achieve complete repression of sfGFP. For the cases where the translation was not completely repressed, the MGA traces display a discernible peak (Fig. 4 D). Our microplate reader was not able to detect MGA signal below 50 nM of mRNA (Supp. Fig. 4 C).

Similar to Csy4, another enhanced CFE extract was produced in which the promoter J23108, driving the expression of dCas9, was replaced by a weaker promoter, J23105 (Supp. Fig. 18 A). Three of the previous UTR constructs were tested along with different concentrations of sg-2 ranging from 2 nM to 10 nM. Corresponding sfGFP fold changes are displayed in Supp. Fig. 18 B. Examples of fluorescence time traces for lrp constructs are shown in Supp. Fig. 19. For rpiV+sD and lrp, we observed a higher dynamic range in the J23108-dCas9 CFE extract while the ion construct displayed a similar fold change in both extracts. This suggests that tuning the ON/OFF level of synthetic circuits could be achieved by combining UTR constructs and enhanced CFE extracts producing different amounts of dCas9. Synthetic biologists could take advantage of this variety of response functions for building customized devices possessing particular signal processing abilities.

### Enhanced CFE extracts for RNA-based circuitry

#### Csy4 increases the dynamic range of a RNA-based AND gate

To apply our enhanced CFE extracts to the prototyping of synthetic circuits, we sought to leverage the RNA-based circuits previously engineered in CFE system. ^21^ Out of several AND-gates based on the small transcriptional activators (STARs) and toehold switches, we selected the sense 6 - toehold 2 (S6T2) gate since its relatively low yield has potential for improvement ^21^. The Csy4 hairpin replaced the linker between the target 6 and toehold 2 components to test if the dynamic range of the gate could be improved by the endoribonuclease (S6-Csy4-T2) (Fig. 5 A). STAR-(activating transcription) and trigger-(activating translation) encoding plasmids were titrated according to the previous characterizations in TXTL (Transcription-Translation) systems ^21^ and dose-response matrices were performed accordingly in both WT and Csy4 CFE extracts. We confirmed previous results in that about three times less trigger plasmid than STAR plasmid was needed to maximize the dynamic range of the gates. The two constructs displayed similar sfGFP dose-responses in the WT CFE extract (Fig. 5 B left). Conversely, S6-Csy4-T2 exhibited higher MGA production rates than S6T2, except at low STAR concentrations (0-3 nM), where low MGA production rates were detected for both constructs. Similar transcriptional results were found in the Csy4 CFE extract (Fig. 5 C). This suggests that S6-Csy4-T2 mRNA may have a higher stability than S6T2. Strikingly, the S6-Csy4-T2 gate displayed up to fourfold increase of sfGFP compared to the S6T2 gate in the Csy4 extract for some combination of STAR/trigger concentrations (Fig. 5 B right). In conclusion, we observed a dramatic enhancement of the translation process of the S6-Csy4-T2 construct expressed in the Csy4 CFE extract. The MGA data suggests this TL enhancement may be decoupled from the high TX rate found with S6-Csy4-T2. In other words, the construct S6-Csy4-T2 displayed an enhancement of the mRNA levels due to the Csy4 hairpin alone, while the translation of sfGFP was only enhanced in the presence of the endoribonuclease Csy4.

**Figure 5:**
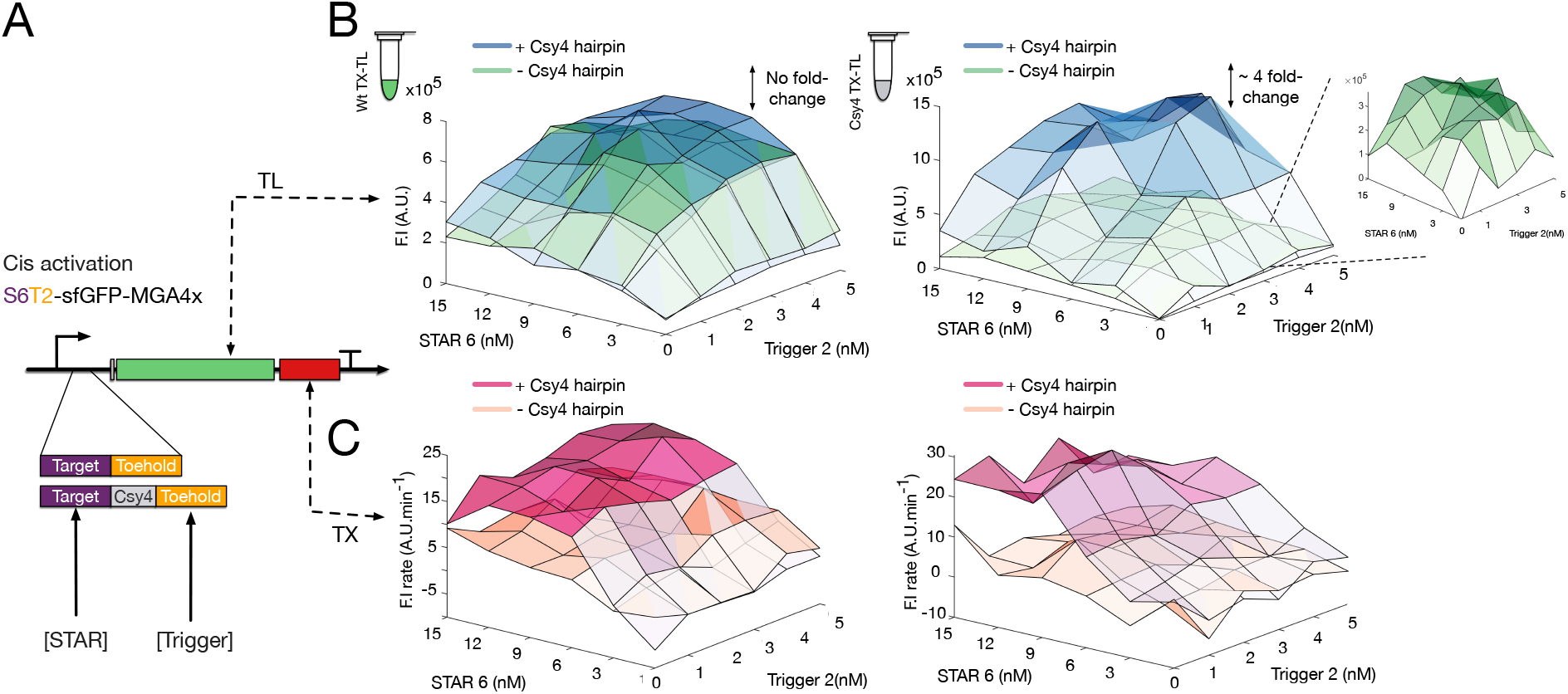
Csy4 activity enhances the dynamic range of S6T2 in the enhanced extract. A) Scheme of the S6T2-sfGFP-MGA4x constructs, with and without the Csy4 hairpin between the sense 6 and toehold 2 components. B) sfGFP endpoint fluorescence measurements of the two gate variants in WT CFE extract (left) and in Csy4 CFE extract (right) with (blue) and without Csy4 hairpin (green). C) Maximal MGA fluorescence production rate of the two gates in WT CFE (left) and in Csy4 CFE (right) with (pink) and without (orange) Csy4 hairpin.

#### sRNA operons can be processed with Csy4 to reduce AND-gate leakage

We next tested if the efficiency of trans-acting small RNA (sRNA) operons could be improved with the use of our enhanced CFE extracts. Csy4 has previously been shown to improve the processing of sRNA operon encoding multiple sgRNAs, by reducing the interference from cis sequences. ^37,38^ We ask here if this paradigm can be extended to sRNA operons encoding a combination of RNA activators and sgRNAs.

We designed two versions of an operon containing three different small RNAs : STAR 6, trigger 3, and sg-2. They were assembled in tandem without any linker in the first version (I1), and in the second version (I2), a Csy4 hairpin was inserted between each component (Fig. 6 A). The two trans-acting operons were tested in the presence of 2 nM of the best performing AND-gate, ^21^ sense 6 - toehold 3 (S6T3), in the WT and enhanced CFE extracts, and normalized to 2 nM of the standard reporter plasmid (to be compared). Fluorescent time traces can be found in Supp. Fig. 21. In the WT and Csy4 CFE extracts, the circuit should act as an AND-logic, with only two inputs used, since dCas9 is absent. The corresponding experiments showed no differences between I1 and I2 on the activation level of the S6T3 gate (Fig. 6 C left). This suggests either that Csy4 does not enhance the processing of this specific sRNA operon, or the sRNA components possess already optimal co-folding structures when expressed together. We titrated the sRNA operons from 1 to 20 nM to detect any saturation-related effect, however, no difference in activation was found for any inputs plasmid concentration (Supp. Fig. 20). The circuit was then tested with the same conditions in the dCas9 CFE extract (Fig. 6 C right). In presence of dCas9, the sgRNA should act as a third input repressing the gate. I1 and I2 exhibited the same level of repression in the dCas9 CFE extract, around 5 times lower than with I1 and I2 in WT and Csy4 CFE extracts. We repeated the repression experiment in a blended CFE extract made of WT and dCas9 extracts in a 1:1 ratio (Fig. 6 B). Similar levels of repression were found as in the dCas9 extract alone. Finally, to test whether Csy4 can reduce the circuit’s leakage, we blended the Csy4 extract with the dCas9 extract in a 1:1 ratio (Fig. 6 B). In this blended CFE extract, I1 showed a similar repression level as in the dCas9 CFE extract (Fig. 6 C right). Conversely, the I2 operon displayed a sfGFP yield similar to the template alone (-I1/-I2), suggesting better processing of sg-2, and therefore, of the gate repression. The reduction of the leakage in the blended CFE extract enabled the increase of the gate’s dynamic range by about 10-fold (in regards to fluorescent endpoints).

**Figure 6:**
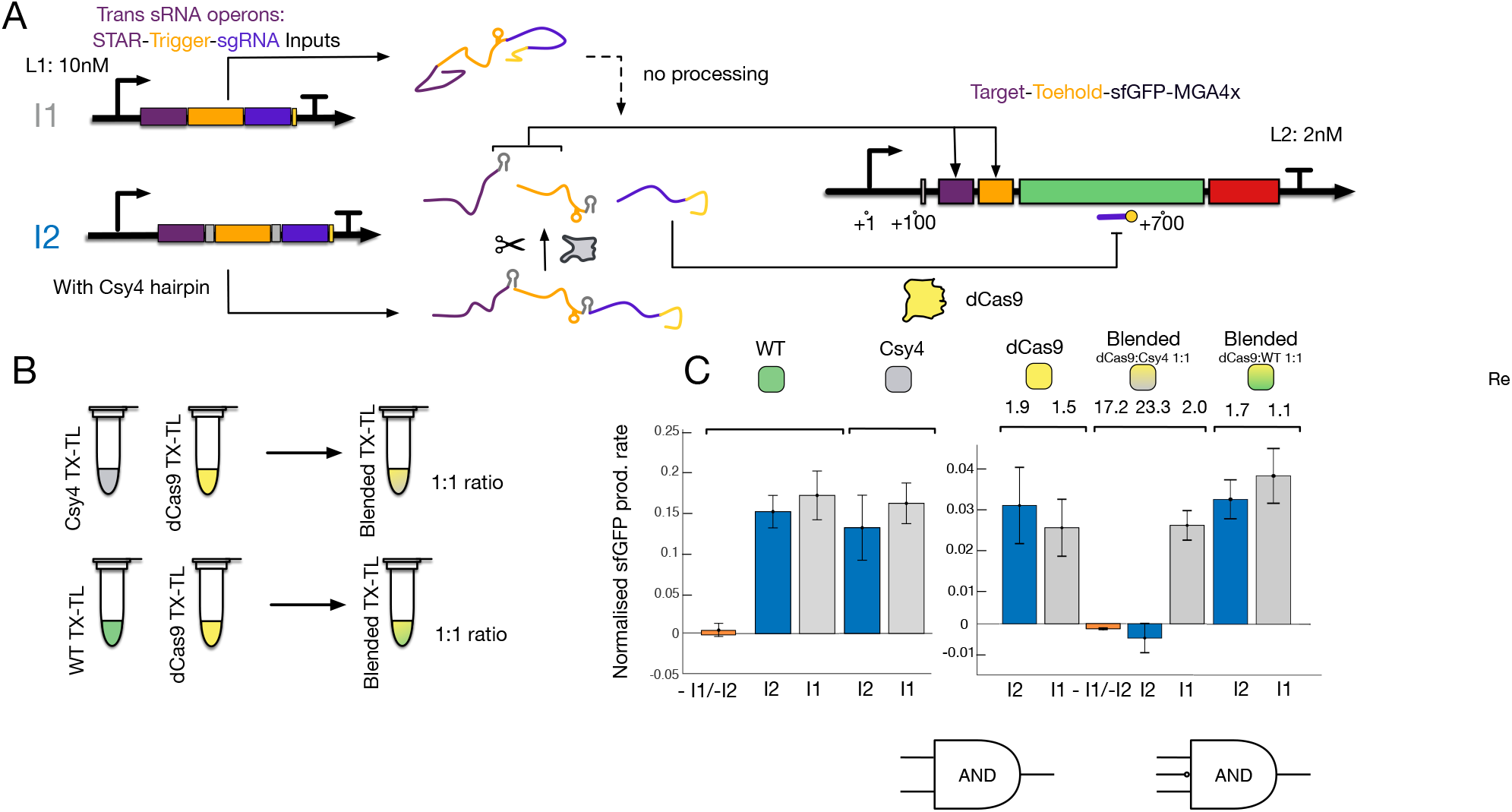
Csy4 and Cas9 CFE extracts can be blended to enhance the functionality of sRNA-operon in synthetic circuits. A) Scheme of the sRNA operon constructs. I2 contains the Csy4 processing hairpin between the STAR, trigger and sgRNA components. STAR 6 and trigger 3 activate the expression of sfGFP, while sg-1 inhibits it. B) CFE extracts can be mixed to form ”blended” CFE extracts (shaded). C) sfGFP production rates for 2 nM of S6T3 gate plasmid (-I1/-I2, orange) and 10 nM of either I1 (grey) or I2 (blue) in four different CFE extracts. Fold-changes are computed from the endpoint fluorescence values. Error bars represent the standard deviation of the mean from three technical replicates.

#### RNA-based NAND gate

Finally, we wanted to explore whether more complex logic circuit prototyping based solely on RNA parts was possible. We sought to build a NAND-gate, which possesses a key property in electrical engineering. Indeed, it is possible to implement any Boolean function based only on NAND-operations. One way to build a NAND-gate is to combine an AND and NOT gate (Fig. 7 A). We used a combination of two orthogonal STARs for the AND module (STAR 1 and STAR 6) ^39^ driving the expression of sg-2, which in turn, should repress the transcription of the targeted sfGFP reporter (Fig. 7 A). To facilitate the processing of the RNA elements placed in tandem, we inserted a Csy4 hairpin between each of them. To assess RNA-based NAND-gate function, we titrated the concentrations of STAR 1 and STAR 6 in presence of 10 nM of Target 6 -Target 1 -sg-2 (T6T1Sg2) plasmid and 2 nM of the reporter plasmid. The corresponding dose-response functions in dCas9 and blended dCas9/Csy4 CFE extracts are shown in Supp. Fig. 22 and Fig. 7 B. The NAND logic was successfully implemented in both CFE extracts, and the transition point from high to low sfGFP expression was shifted to lower STAR concentrations in the blended CFE extract. The maximum fold-change between ON-state and the lowest OFF-state also increased from 4.3-to 6.7-fold in the blended CFE extract. We attributed the overall sharper transition and increased dynamic range of the NAND gate in the blended extract to the cleavage of the STAR targets and sg-2 components by Csy4. However, we observed a decrease in the sfGFP production rate at increasing STAR concentrations, suggesting that STAR 1 and STAR 6 lack orthogonality for their sense target at high concentrations. The cause of this leakage could originate from the design of the STAR targets, which share a common terminator sequence. The STAR activators, in turn, also share a small sequence complementary to the terminator stem. 39 We are confident that the design of other STARs based on a new set of terminators would enable the engineering of more robust NAND gates.

**Figure 7:**
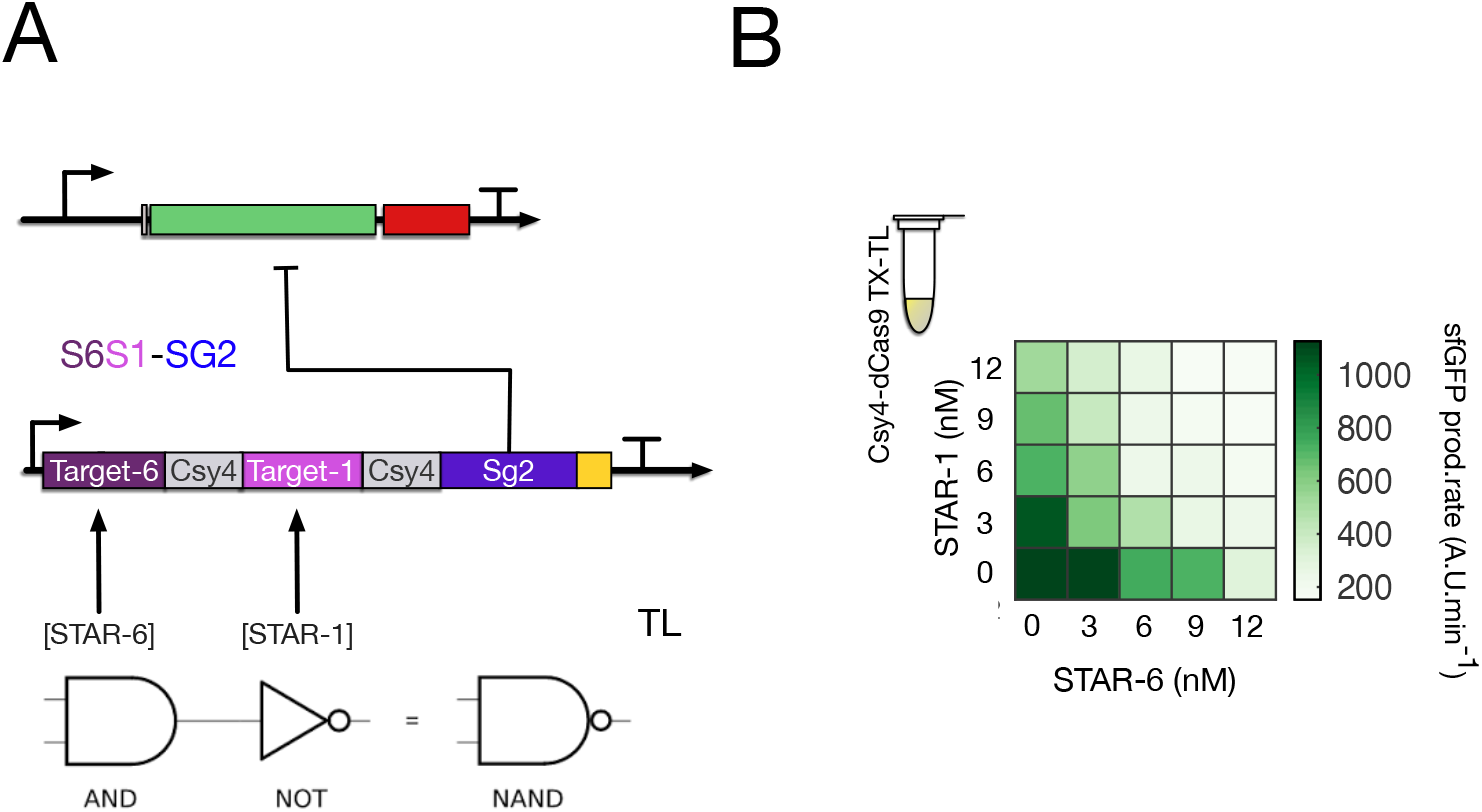
Enhanced CFE extracts can be used for NAND logic prototyping. A) Scheme of the NAND circuit based on the coupling of an AND and NOT gates. Two orthogonal STARs activate the transcription of sg2, which in turn inhibits sfGFP production. B) Heatmap of the sfGFP production rates for the NAND gate titrated with STAR-1 and STAR-6 encoding plasmids in blended Csy4-dCas9 extract. Other parts concentrations were set to 2 nM (reporter plasmid) and 10 nM (sg-2 expressing plasmid).

### Conclusion

In this work, we demonstrated that the recent cell-free extract protocol from Silverman et al. ^6^ can be easily adapted to produce augmented CFE extracts. In particular, we showed that pre-expressing CRISPR-associated proteins dCas9 and Csy4 does not diminish the final mRNA or protein yield compared to the WT CFE extract. We characterized four highly functional sgRNAs as well as different 5’ UTR constructs which could serve as a toolbox of parts for our augmented CFE extracts. Csy4 cleavage activity displayed high efficiency for DNA concentrations traditionally used in cell-free circuit prototyping. We anticipate this could have implications in the implementation of Csy4 derivatives in cell-free systems, such as the inactivated Csy4^40^ or Csy4-mediated RNA switches. ^34^ The concentration of CRISPR-associated proteins can be tuned via the promoter strength of the expression cassette to achieve various processing efficiencies. The use of MGA aptamers enabled us to monitor the mRNA dynamics in diverse circuitry configurations, constituting novel examples of using a dual-reporter system. This will be helpful to parameterize computational models, in particular, to enhance RNA-based models whose transcriptional kinetics was usually based only on inference. ^22,41^

Additionally, we used the enhanced CFE extracts to improve and design RNA-based circuitry. The use of dCas9 and the associated sgRNAs enable diversification of Boolean logic functions from the previously characterized RNA-based components.^19,21^ We demonstrated that Csy4 could reduce leakage of trans-acting sRNA operons, as well as increase the dynamic range of a previously engineered RNA-based AND gate. Taking advantage of the dual reporter system for the multi-level regulated gate, we were able to conclude that Csy4 processing specifically enhances the translational activity. This could encourage the use of the short cleaving hairpin in RNA-based circuits or offer the possibility to take advantage of existing compatible DNA parts.^39,42^ Future work could extend the one-pot strategy to include additional enhanced CFE extracts. For instance, producing RNA chaperones such as Hfq could increase the binding affinities of the small RNAs and stabilize target mRNAs to increase the efficiency of RNA-based circuits. ^43^ Finally, we hope this enhanced CFE platform will facilitate access to cell-free circuit prototyping tools, in areas ranging from point-of-care diagnostics to educational projects.

## Experimental

### Cell-free extract production

*E. coli* BL21 Rosetta 2 were streaked overnight on an agar plate containing chloramphenicol (and ampicillin, for the production of enhanced extracts). One colony was picked and inoculated overnight in 50 mL 2xYT supplemented with antibiotic(s) for growth at 37°C. After a minimum of 15 hours, 10 to 20 mL of the stationary culture was used to inoculate 200 mL to 1 L of 2xYT + P media (16 g/L tryptone, 10 g/L yeast extract, 5 g/L sodium chloride, 7 g/L potassium phosphate dibasic, 3 g/L potassium phosphate monobasic) in a 1 L to 5 L baffled flask. For the experiments not related to the study of optical density (*OD*_600_), cells were grown at 37 °C and 200 rpm to 3.5 ± 0.1 *OD*_600_. Centrifuge bottles were filled up to 300 mL and centrifuged for 10 minutes at 4000 x g at 4 °C and supernatants were discarded. The pellets were washed three times with 25 mL buffer S30A (50 mM Tris-base, 14 mM Mg-glutamate, 60 mM K-glutamate, 2 mM DTT, brought to pH 7.7 with acetic acid). Each wash step was followed by a centrifugation step at 4000 g at 4 °C for 10 minutes. A fourth centrifuging step at 3000 g at 4°C for 10 minutes enabled the removal of the remaining traces of buffer. The pellets were then resuspended in 1 mL of Buffer S30A per gram of dry pellet. 700 μL of the suspension was split into the necessary number of 2 mL Eppendorf tubes. The pellet suspensions were then lysed with a sonicator (QSonica Q125 with a 3.175 mm diameter probe, 50% amplitude, 20 kHz, and 10 seconds ON/OFF pulses). Each sample was sonicated until reaching 500 J unless otherwise indicated. 1 mM of DTT was added to each crude lysate immediately after sonication. The cell lysate was then centrifuged for 10 minutes at 4 °C and 12000 g. The supernatant was removed and placed into an incubator set up at 37 °C and 200 rpm for 80 minutes. After this run-off reaction, the supernatant was centrifuged again for 10 minutes at 4 °C and 12000 g. The supernatant was aliquoted as 40 μL samples into PCR tubes. The aliquots were snap-frozen into liquid nitrogen and stored at −80 °C.

### Cell-free experiment

The final CFE reaction mixture is composed of the following reagents: 10 mM ammonium glutamate; 1.2 mM ATP; 0.850 mM each of GTP, UTP, and CTP; 0.034 mg/mL folinic acid; 0.175 mg/mL yeast tRNA; 2 mM amino acids; 30 mM 3-PGA; 0.33 mM NAD; 0.27 mM CoA; 1 mM putrescine; 1.5 mM spermidine; 57 mM HEPES. The standard concentration of Mg-glutamate and K-glutamate were kept respectively at 10 mM and 140 mM. Some experiments involved the titration of K-glutamate between 80 mM and 200 mM. Before each use, the reactions were thawed at 4 °C and RNase inhibitor (MOLO) was added to the mix (0.01). For experiments involving MGA read-out, the plates were chilled beforehand in the −20 °C freezer to slow the start of the kinetic reactions. Reactions were prepared either to the 2 or 5 μL scale in triplicate. For 2 μl reactions, the master mix was dispensed into a PCR 384 well plate (Greiner) with the help of an automatic dispenser (Eppendorf Repeater E3). The volume was then brought up with plasmid DNA constructs containing the genetic circuits and nuclease-free water (0.5) by using either an I-Dot One SC (Dispendix), or by pipetting. The microplate was then quickly centrifuged and covered with a transparent sealer (Ampliseal, Greiner). Finally, the microplate was vortexed and centrifuged again before being ready for either kinetic or endpoint measurements.

#### Incubation and kinetic measurements

PCR 384-well white plates containing assembled cell-free reactions and DNA were measured with a ClarioSTAR (BMG Biotech). The temperature was set to 29 °C for the duration of the experiment (10 hours). The sfGFP and MGA signals were measured every 2 minutes with the following wavelengths: 485/520 (Ex/Em) and 610/650 (Ex/Em), with the enhanced dynamic gain range enabled and z-height optimised according to each experiment. For all experiments, the negative control with no DNA template was prepared in triplicate and the corresponding kinetics or end-points were subtracted from the reported fluorescence values. For experiments measuring the output of circuits based on the expression of sRNA repressors or activators, the same amount of plasmids encoding for decoy sRNAs was added to the positive control condition.

For experiments where we needed to compare data from different enhanced CFE extracts, the fluorescence values were normalised to a positive control condition containing the same concentration of Pr-sfGFP-MGA4x plasmid expressed in triplicates in each CFE extract.

#### Concentrations of DNA parts

Except for Supp. Fig. 14, all RNA parts were co-expressed in cell-free reactions from DNA templates. For the AND gates in Fig. 5 and Fig. 6 as well as the NAND gate is Fig. 7, the reporter plasmid was set to 2 nM. Except for the dose-reponse experiments or stated otherwise, the concentration of STAR/trigger/sg RNA constructs were set to 10 nM.

### Assembly of expression constructs

The construction of the plasmid library is based on plasmids previously published: pJBL2801 was a gift from Julius Lucks (Addgene plasmid 71207) and was used for cloning the UTR constructs, the sgRNAs, and the dCas9/Csy4/mRFP1 production plasmids; pJBL2807 was a gift from Julius Lucks (Addgene plasmid 71203) and was used for cloning the sense, toehold and the nine combined gates; pET21b-RL015A was a gift from Sheref Mansy (Addgene plasmid 42134) and used for the construction of J23119-GFP-mCherry plasmid; pJ1996 v2 plasmid was a gift from Yolanda Schaerli (Addgene plasmid 140664) and used for co-expression of Csy4 for *in vivo* experiments. Golden Gate and Gibson assembly were used for cloning the inserts with length longer than 100 bps. Q5 site directed mutagenesis kit from NEB was used for cloning the inserts smaller than 100 bps. Oligos used for the various assemblies or mutagenesis were ordered from Sigma Aldrich. The sRNA operons and the tandem repeat of MGAs were ordered from Genscript. Sanger sequencing verified all constructed plasmids. Plasmids were prepared using a NucleoBond Extra Midi Plus (Macherey-Nagel) and followed by isopropanol precipitation and eluted with 10 mM Tris at pH=8.5 and quantified using a NanoDrop 1000. Those having an insufficient purity were subject to a Phenol-chloroform extraction followed by a novel step of isopropanol precipitation. The sequence of each part used is available in the supplemental information.

### *In vivo* Experiment

*E. coli* colonies (NEB 5-alpha) containing GFP-mCherry or GFP-Csy4-mCherry with and without pJ1996 v2 plasmids were selected and inoculated in 2 mL of LB containing selective antibiotics and grown for 6 h at 37 °C, 200 rpm. The cultures were centrifuged and resuspended in Hi-DEF Azure Media (Teknova) with 4% glycerol. 120 μL of 0.05 *OD*_600_ bacterial cultures were added per well on a 96 well Sensoplate Microplate (Greiner Bio One, Austria), and incubated at 37 °C, 200 rpm for 16 h. Fluorescence was determined using a ClarioSTAR (BMG Biotech). Fluorescence values were corrected using blank samples and normalised by *OD*_600_ values.

### Gel electrophoresis

CFE extracts were diluted to 1:5, 1:10 and 1:20 in Buffer A (50 mM Tris-HCl, 14 mM Mg-glutamate, 60 mM K-glutamate, pH 7.7). BlueStar Prestained Protein Marker (NIPPON Genetics Europe, Germany) was used. For positive control and quantification of dCas9 in CFE extract, commercially purified dCas9 protein was used (Sigma Aldrich, US). Tris-glycine SDS running buffer (25 mM Tris, 250 mM glycine, 0.1% SDS) was used for electrophoresis. After electrophoresis, gels (4–20% Mini-PROTEAN^®^ TGX™ Precast Protein Gels, Bio-Rad, US) and Tris-glycine SDS running buffer (25 mM Tris, 250 mM glycine, 0.1% SDS) was used for electrophoresis.” were stained with Coomassie Blue G-250 (ThermoScientific, US) for the rapid detection of protein bands. The bands were detected by E-Box (Vilber, Germany).

### Western Blotting

After electrophoresis, proteins were transferred onto the PVDF membrane from Trans-Blot Turbo Mini 0.2 μm PVDF Transfer Pack (Bio-Rad, US) at 2.5 A for 3 min by using Trans-Blot Turbo Transfer System (Bio-Rad, US). The blotted PVDF membranes were directly blocked with an EveryBlot Blocking Buffer (Bio-Rad, US) for 5 min and then probed with primary antibodies, anti-Cas9 (7A9-3A3 Mouse mAb, Cell Signalling, US) or anti-FLAG (Monoclonal ANTI-FLAG^®^ M2 antibody produced in mouse, Sigma Aldrich, US), overnight at 4°C. After washing with TBST (TBST-10X, Cell Signalling, US) 3 times, the membrane was probed with a secondary antibody (anti-Mouse Rat IgG, Invitrogen, US) for 1 h at room temperature. The immunoreactive bands were detected by the use of Clarity Western ECL Substrate (Bio-Rad, US) on Amersham Imager 600 (GE Healthcare, US). The protein quantification was performed with the use of ImageJ 1.53a (Wayne Rasband National Institutes of Health, US).

## Acknowledgement

This work was supported by the Landesoffensive für wissenschaftliche Exzellenz as part of the LOEWE Schwerpunkt CompuGene. HK acknowledges support from the European Research Council (ERC) with the consolidator grant CONSYN (grant no. 773196).

## References

(1) Silverman, A. D.; Karim, A. S.; Jewett, M. C. Cell-free gene expression: an expanded repertoire of applications. Nature Reviews Genetics 2020, 21, 151–170.

(2) Laohakunakorn, N.; Grasemann, L.; Lavickova, B.; Michielin, G.; Shahein, A.; Swank, Z.; Maerkl, S. J. Bottom-Up Construction of Complex Biomolecular Systems With Cell-Free Synthetic Biology. Frontiers in Bioengineering and Biotechnology 2020, 8, 213.

(3) Sun, Z. Z.; Hayes, C. A.; Shin, J.; Caschera, F.; Murray, R. M.; Noireaux, V. Protocols for Implementing an Escherichia coli Based TX-TL Cell-Free Expression System for Synthetic Biology. Journal of Visualized Experiments 2013, e50762.

(4) Takahashi, M. K.; Hayes, C. A.; Chappell, J.; Sun, Z. Z.; Murray, R. M.; Noireaux, V.; Lucks, J. B. Characterizing and prototyping genetic networks with cell-free transcription-translation reactions. Methods 2015, 86, 60–72.

(5) Lian, Q.; Cao, H.; Wang, F. The Cost-Efficiency Realization in the Escherichia coli-Based Cell-Free Protein Synthesis Systems. Applied Biochemistry and Biotechnology 2014, 2351–2367.

(6) Silverman, A. D.; Kelley-Loughnane, N.; Lucks, J. B.; Jewett, M. C. Deconstructing cell-free extract preparation for in vitro activation of transcriptional genetic circuitry.

(7) Chappell, J.; Jensen, K.; Freemont, P. S. Validation of an entirely in vitro approach for rapid prototyping of DNA regulatory elements for synthetic biology. Nucleic Acids Research 2013, 41, 3471–3481.

(8) Siegal-gaskins, D.; Noireaux, V.; Murray, R. M. Biomolecular resource utilization in elementary cell-free gene circuits. American Control Conference (ACC) 2013, 1531–1536.

(9) Nagaraj, V. H.; Greene, J. M.; Sengupta, A. M.; Sontag, E. D. Translation inhibition and resource balance in the TX-TL cell-free gene expression system. Synthetic Biology 2017, 2, DOI: 0.1093/synbio/ysx005.

(10) Moore, S. J.; MacDonald, J. T.; Wienecke, S.; Ishwarbhai, A.; Tsipa, A.; Aw, R.; Kylilis, N.; Bell, D. J.; McClymont, D. W.; Jensen, K.; Polizzi, K. M.; Biedendieck, R.; Freemont, P. S. Rapid acquisition and model-based analysis of cell-free transcription–translation reactions from nonmodel bacteria. Proceedings of the National Academy of Sciences 2018, 115, E4340–E4349.

(11) Matsubayashi, H.; Ueda, T. Cell-free protein synthesis system. Kobunshi 2014, 63, 379–381.

(12) Garamella, J.; Marshall, R.; Rustad, M.; Noireaux, V. The All E. coli TX-TL Toolbox 2.0: A Platform for Cell-Free Synthetic Biology. ACS Synthetic Biology 2016, 5, 344–355.

(13) Dopp, J. L.; Jo, Y. R.; Reuel, N. F. Methods to reduce variability in E. Coli-based cell-free protein expression experiments. Synthetic and Systems Biotechnology 2019, 4, 204–211.

(14) Guan, D.; Chen, Z. Challenges and recent advances in affinity purification of tag-free proteins. Biotechnology Letters 2014, 36, 1391–1406.

(15) Soye, B. J. D.; Gerbasi, V. R.; Thomas, P. M.; Kelleher, N. L.; Jewett, M. C. A Highly Productive, One-Pot Cell-Free Protein Synthesis Platform Based on Genomically Re-coded Escherichia coli. Cell Chemical Biology 2019, 26, 1743–1754.e9.

(16) Dudley, Q. M.; Anderson, K. C.; Jewett, M. C. Cell-Free Mixing of Escherichia coli Crude Extracts to Prototype and Rationally Engineer High-Titer Mevalonate Synthesis. ACS Synthetic Biology 2016, 5, 1578–1588.

(17) Karim, A. S.; Jewett, M. C. A cell-free framework for rapid biosynthetic pathway prototyping and enzyme discovery. Metabolic Engineering 2016, 36, 116–126.

(18) Jaroentomeechai, T.; Stark, J. C.; Natarajan, A.; Glasscock, C. J.; Yates, L. E.; Hsu, K. J.; Mrksich, M.; Jewett, M. C.; DeLisa, M. P. Single-pot glycoprotein biosynthesis using a cell-free transcription-translation system enriched with glycosylation machinery. Nature Communications 2018, 9, 2686.

(19) Marshall, R. et al. Rapid and Scalable Characterization of CRISPR Technologies Using an E. coli Cell-Free Transcription-Translation System. Molecular Cell 2018, 69, 146–157.e3.

(20) Liao, C.; Ttofali, F.; Slotkowski, R. A.; Denny, S. R.; Cecil, T. D.; Leenay, R. T.; Keung, A. J.; Beisel, C. L. Modular one-pot assembly of CRISPR arrays enables library generation and reveals factors influencing crRNA biogenesis. Nature Communications 2019, 10, 2948.

(21) Lehr, F.-X.; Hanst, M.; Vogel, M.; Kremer, J.; Göringer, H. U.; Suess, B.; Koeppl, H. Cell-Free Prototyping of AND-Logic Gates Based on Heterogeneous RNA Activators. ACS Synthetic Biology 2019, 8, 2163–2173.

(22) Hu, C. Y.; Takahashi, M. K. Engineering a Functional Small RNA Negative Autoregulation Network with Model-Guided Design. ACS Synthetic Biology 2018, 7, 1507–1518.

(23) Collias, D.; Marshall, R.; Collins, S. P.; Beisel, C. L.; Noireaux, V. An educational module to explore CRISPR technologies with a cell-free transcription-translation system. Synthetic Biology 2019, 4, ysz005.

(24) Kwon, Y.-C.; Jewett, M. C. High-throughput preparation methods of crude extract for robust cell-free protein synthesis. Scientific Reports 2015, 5, 8663–8670.

(25) Cole, S. D.; Beabout, K.; Turner, K. B.; Smith, Z. K.; Funk, V. L.; Harbaugh, S. V.; Liem, A. T.; Roth, P. A.; Geier, B. A.; Emanuel, P. A.; Walper, S. A.; Chávez, J. L.; Lux, M. W. Quantification of Interlaboratory Cell-Free Protein Synthesis Variability. ACS Synthetic Biology 2019, 2080–2091.

(26) Sun, Z. Z.; Hayes, C. A.; Shin, J.; Caschera, F.; Murray, R. M.; Noireaux, V. Protocols for implementing an Escherichia coli based TX-TL cell-free expression system for synthetic biology. Journal of visualized experiments : JoVE 2013, e50762–e50762.

(27) Failmezger, J.; Rauter, M.; Nitschel, R.; Kraml, M.; Siemann-Herzberg, M. Cell-free protein synthesis from non-growing, stressed Escherichia coli. Scientific Reports 2017, 7, 16524.

(28) Yerramilli, V. S.; Kim, K. H. Labeling RNAs in Live Cells Using Malachite Green Aptamer Scaffolds as Fluorescent Probes. ACS Synthetic Biology 2018, 7, 758–766.

(29) Qi, L.; Haurwitz, R. E.; Shao, W.; Doudna, J. A.; Arkin, A. P. RNA processing enables predictable programming of gene expression. Nature Biotechnology 2012, 30, 1002–1006.

(30) Anderson promoter collection. Available: http://parts.igem.org/Promoters/Catalog/Anderson.

(31) Rosano, G. L.; Ceccarelli, E. A. Recombinant protein expression in Escherichia coli: advances and challenges. Frontiers in microbiology 2014, 5, 172–172.

(32) Sternberg, S. H.; Haurwitz, R. E.; Doudna, J. A. Mechanism of substrate selection by a highly specific CRISPR endoribonuclease. RNA (New York, N.Y.) 2012, 18, 661–672.

(33) Marshall, R.; Noireaux, V. Quantitative modeling of transcription and translation of an all-E. coli cell-free system. Scientific Reports 2019, 9, 11980.

(34) Guo, H.; Song, X.; Lindner, A. B. Anti-CRISPR RNAs: designing universal riboregulators with deep learning of Csy4-mediated RNA processing. bioRxiv 2020,

(35) Hu, C. Y.; Varner, J. D.; Lucks, J. B. Generating Effective Models and Parameters for RNA Genetic Circuits. ACS Synthetic Biology 2015, 4, 914–926.

(36) Doench, J. G.; Fusi, N.; Sullender, M.; Hegde, M.; Vaimberg, E. W.; Donovan, K. F.; Smith, I.; Tothova, Z.; Wilen, C.; Orchard, R.; Virgin, H. W.; Listgarten, J.; Root, D. E. Optimized sgRNA design to maximize activity and minimize off-target effects of CRISPR-Cas9. Nature Biotechnology 2016, 34, 184–191.

(37) Nissim, L.; Perli, S.; Fridkin, A.; Perez-Pinera, P.; Lu, T. Multiplexed and Programmable Regulation of Gene Networks with an Integrated RNA and CRISPR/Cas Toolkit in Human Cells. Molecular Cell 2014, 54, 698–710.

(38) Tsai, S. Q.; Wyvekens, N.; Khayter, C.; Foden, J. A.; Thapar, V.; Reyon, D.; Goodwin, M. J.; Aryee, M. J.; Joung, J. K. Dimeric CRISPR RNA-guided FokI nucleases for highly specific genome editing. Nature Biotechnology 2014, 32, 569–576.

(39) Chappell, J.; Westbrook, A.; Verosloff, M.; Lucks, J. B. Computational design of small transcription activating RNAs for versatile and dynamic gene regulation. Nature Communications 2017, 8, DOI: 10.1038/s41467-017-01082-6.

(40) Lee, H. Y.; Haurwitz, R. E.; Apffel, A.; Zhou, K.; Smart, B.; Wenger, C. D.; Laderman, S.; Bruhn, L.; Doudna, J. A. RNA–protein analysis using a conditional CRISPR nuclease. Proceedings of the National Academy of Sciences 2013, 110, 5416–5421.

(41) Hu, C. Y.; Varner, J. D.; Lucks, J. B. Generating effective models and parameters for RNA genetic circuits. bioRxiv 2015,

(42) Santos-Moreno, J.; Tasiudi, E.; Stelling, J.; Schaerli, Y. Multistable and dynamic CRISPRi-based synthetic circuits. Nature Communications 2020, 11, 2746.

(43) Soper, T.; Mandin, P.; Majdalani, N.; Gottesman, S.; Woodson, S. A. Positive regulation by small RNAs and the role of Hfq. Proceedings of the National Academy of Sciences of the United States of America 2010, 107, 9602–9607.

