## Supplementary information for "Functionalizing cell-free systems with CRISPR-associated proteins: Application to RNA-based circuit engineering"

### Supporting Information Available

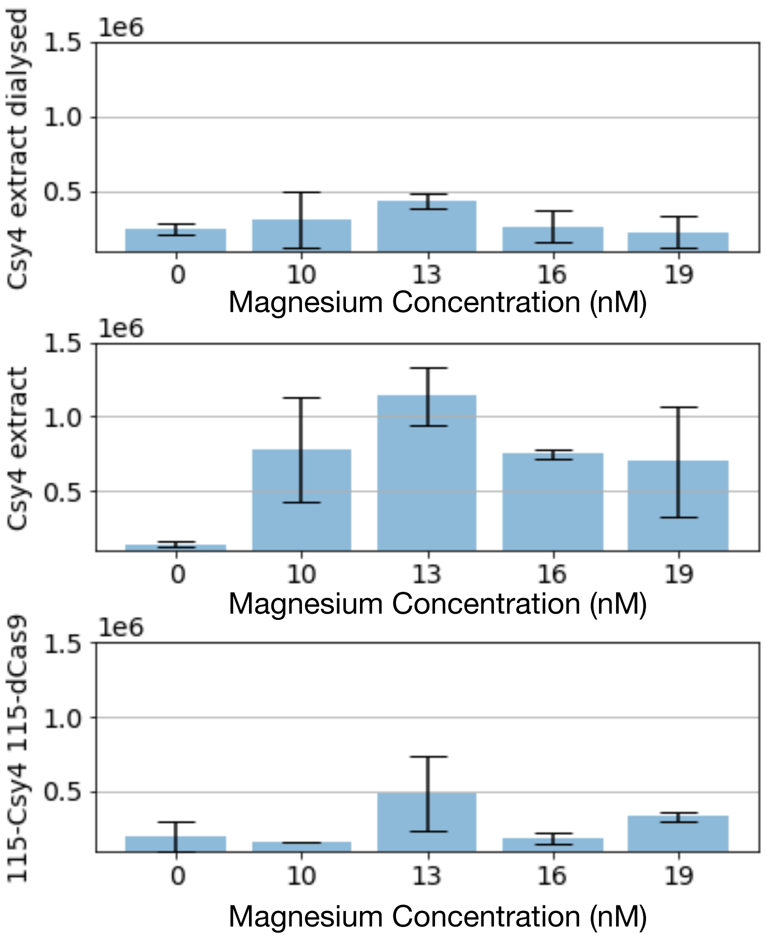


Figure 1: Effect of dialysis and magnesium concentration on CFE extract performance. From top to bottom, J23108-Csy4 CFE extract (with dialysis), J23108-Csy4 CFE extract (without dialysis), J123115-Csy4-J23115-dCas CFE extract (without dialysis). Each condition was tested for 1 nM of reporter plasmid and reported as GFP F.I. (A.U.)

**A. Growth Curves of WT and mRFP**

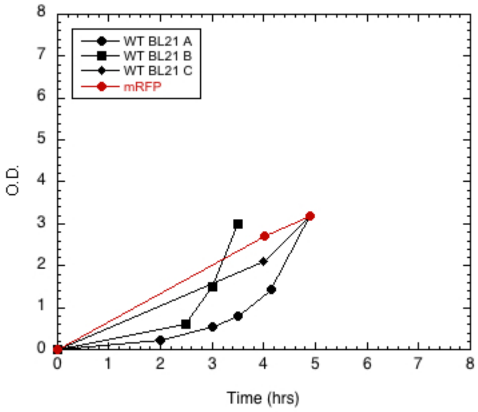


**C. Growth Curves of Csy4**

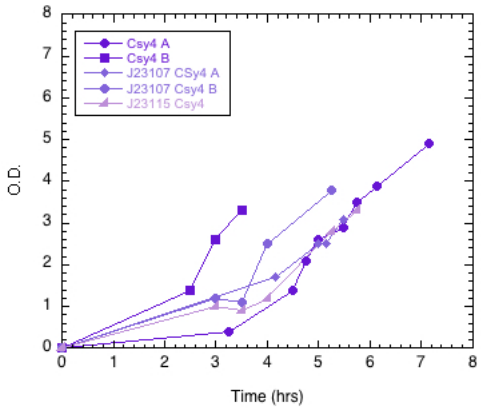


**B. Growth Curves of Cas9**

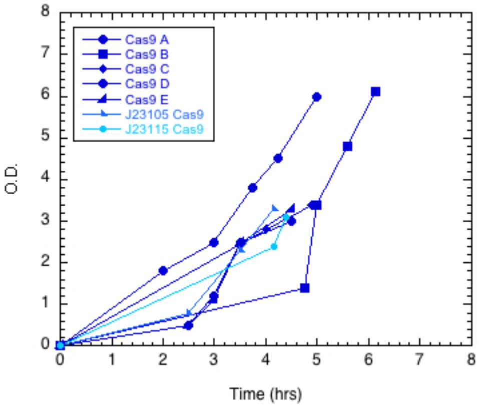


**D. Doubling Time is similar among Extracts**

| Cell Extract | Growth Rate (Rate/Hr) |
|---|---|
| WT BL21 A | 0.24 |
| WT BL21 B | 0.31 |
| WT BL21 C | 024 |
| mRFP | 0.24 |
| Cas9 A | 0.36 |
| Cas9 B | 0.24 |
| Cas9 C | 0.25 |
| Cas9 D | 0.24 |
| Cas9 E | 0.27 |
| J23105 Cas9 | 0.29 |
| J23115 Cas9 | 0.26 |
| Csy4 A | 0.22 |
| Csy4 B | 0.34 |
| J23107 Csy4 | 0.21 |
| J23107 Csy4 | 0.25 |
| J23115 Csy4 | 0.21 |

Figure 2: Growth curves and doubling rates for all extracts. A) Growth curves for WT BL21 and mRFP extracts. B) Growth Curves for Cas9 CFE extracts. C) Growth Curves for Csy4 CFE extracts. D) Table of doubling rates in hours for all CFE extracts. Calculated as $Ln(OD2 - OD1)/Time2 - Time1$.

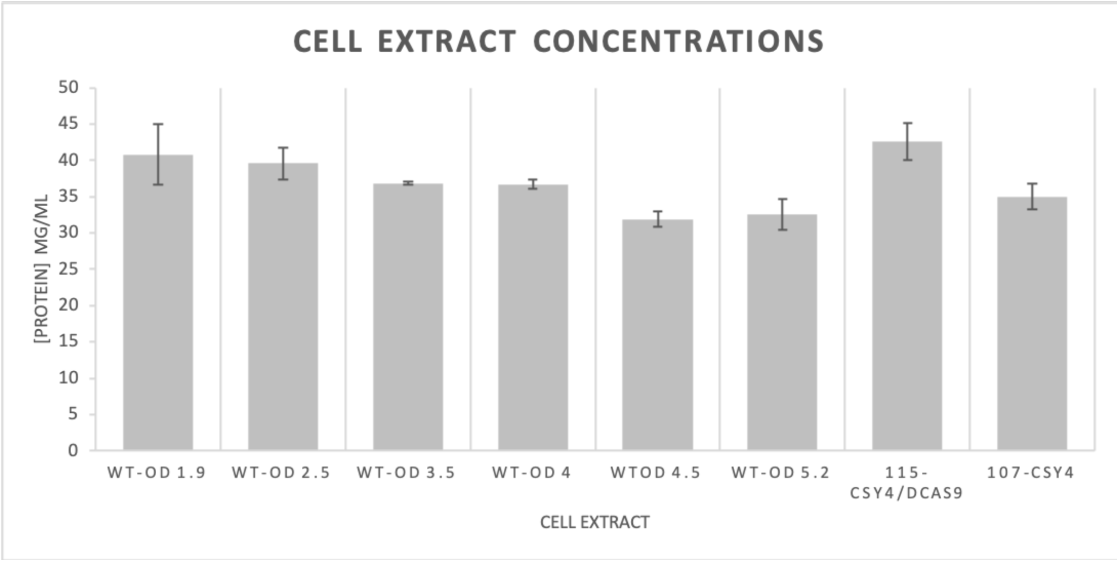


Figure 3: Protein concentration in CFE extracts determined by BSA titration. Error bars represent the standard variation of three technical replicates.

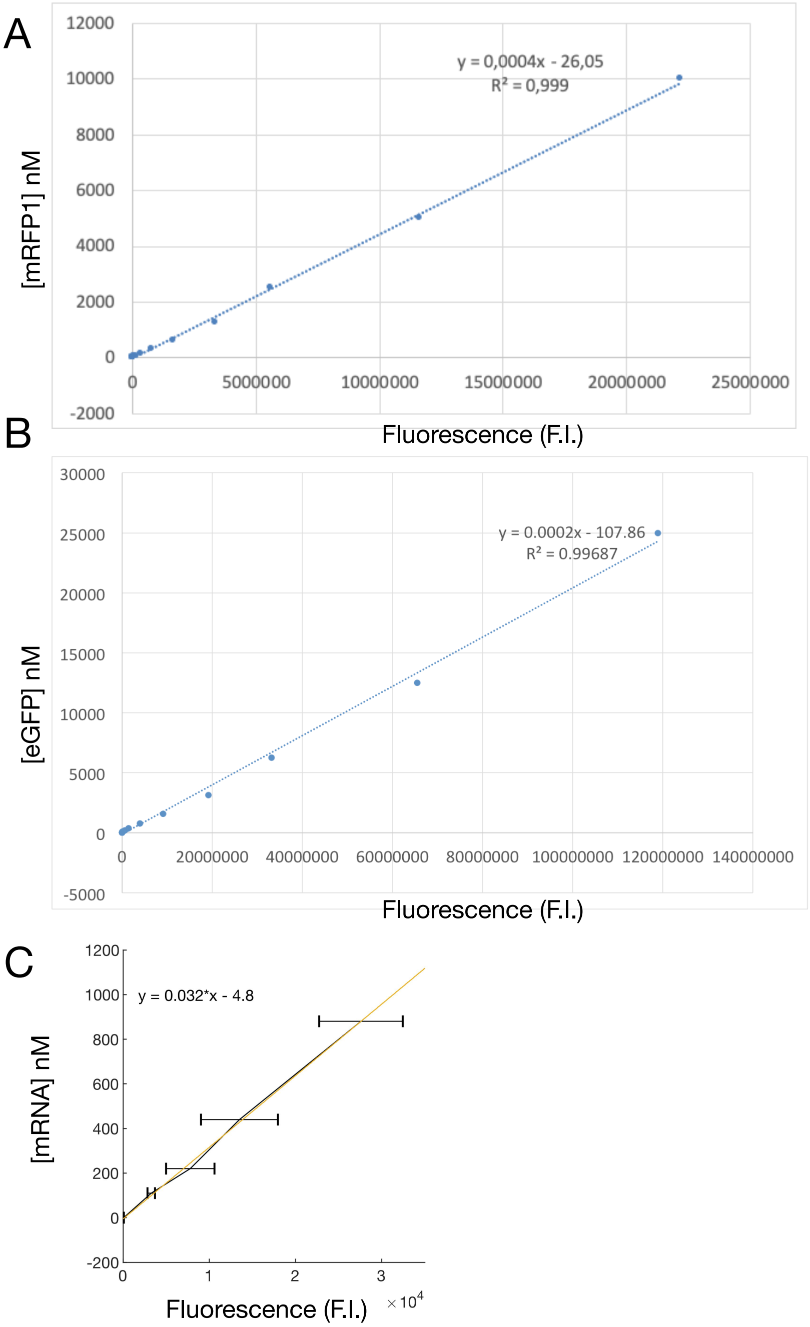


Figure 4: Calibration curves for quantifying protein and mRNA levels in the cell extracts. A) Calibration curve with purified mRFP1. B) Calibration curve with purified eGFP. C) Calibration curve for purified sfGFP-MGA4x mRNA (in CFE energy buffer and 40 nM of MGA dye). A linear fit has been applied to the three calibration curves (Matlab 2019a).

A

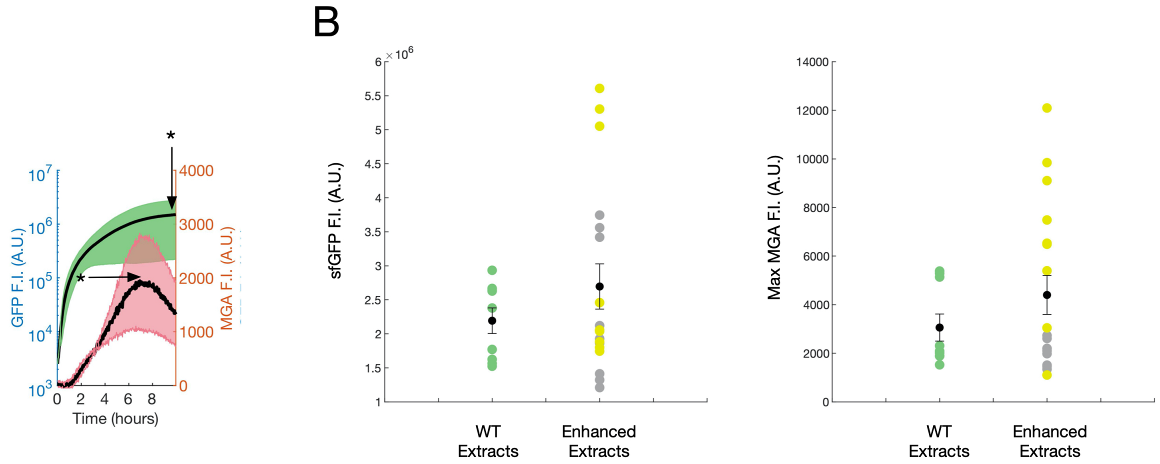


C

| | Mean WT | Mean Enhanced | Std. error WT | Std. error Enhanced | 95% CI lower | 95% CI upper | t | df | p-value |
|---|---|---|---|---|---|---|---|---|---|
| Endpoint GFP | 2,19E+06 | 2,70E+06 | 1,88E+05 | 3,32E+05 | 1.0E+05*-1,517 | 1.0E+05*-3.8278 | 1,0217 | 25 | 0.3167 |
| Max MGA | 3,06E+04 | 4,40E+04 | 5,58E+03 | 8,01E+03 | 1.0E+03*-1,517 | 1.0E+03*-1,1146 | 1,1114 | 25 | 0.2770 |

Figure 5: Yield variation in different batches of WT and enhanced CFE extracts. A) Example of fluorescent time trace for Csy4 CFE extract. The * represent sfGFP endpoints and MGA maximum fluorescence values chosen for the statistical analysis. B) Three batches of WT extract (green) and six batches of enhanced CFE extracts (yellow for dCas9 and grey for Csy4) were tested with 5 nM of standard reporter plasmid (Pr-sfGFP-MGA4x) and three replicates were performed for each extract. Scatter plots for sfGFP (left) and MGA (right) are shown accordingly. The black dots represent the mean and the error bars correspond to the standard error of the mean. C) Summary result of two-samples t-tests performed for sfGFP endpoint and MGA maximal fluorescence. In either case, the t-test does not reject the null hypothesis at the default 5 % significance level.

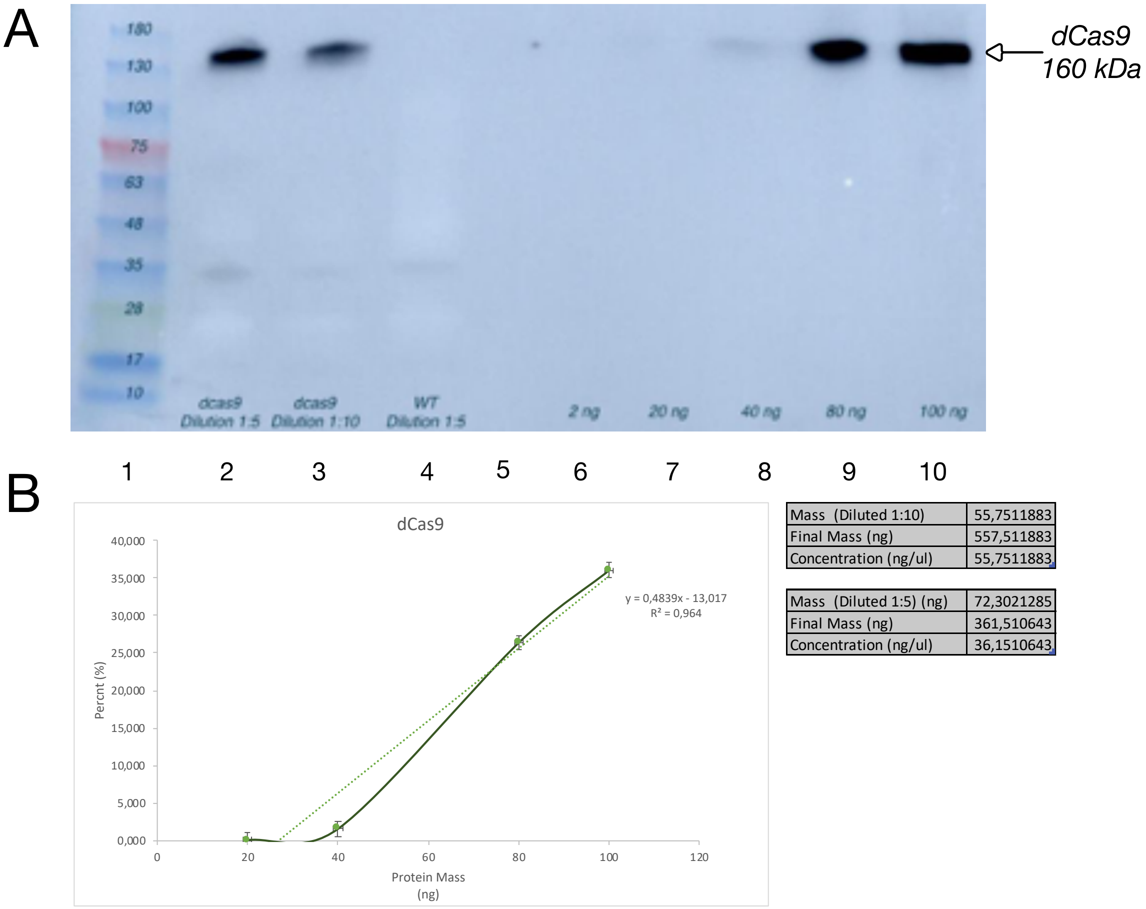


| | |
|---|---|
| Mass (Diluted 1:10) | 55,7511883 |
| Final Mass (ng) | 557,511883 |
| Concentration (ng/ul) | 55,7511883 |

| | |
|---|---|
| Mass (Diluted 1:5) (ng) | 72,3021285 |
| Final Mass (ng) | 361,510643 |
| Concentration (ng/ul) | 36,1510643 |

Figure 6: Quantification of dCas9 in enhanced CFE extract. A) dCas9 lysate samples (10 µL) were analysed by 4-20% SDS-PAGE, and the separated proteins were transferred onto PVDF membranes probed with anti-Cas9 antibody. Line 1: ladder. Line 2: dCas9 lysate at 1:5 dilution. Line 3: dCas9 lysate at 1:10 dilution. Line 4: WT lysate at 1:5 dilution. Line 5: empty. Line 6: Cas9 positive control sample of 2 ng. Line 7: Cas9 positive control sample of 20 ng. Line 8: Cas9 positive control sample of 40 ng. Line 9: Cas9 positive control sample of 80 ng. Line 10: Cas9 positive control sample of 100 ng. B) Standard curve obtained from the quantification in A. The amount of targeted dCas9 in the two diluted samples are displayed on the right.

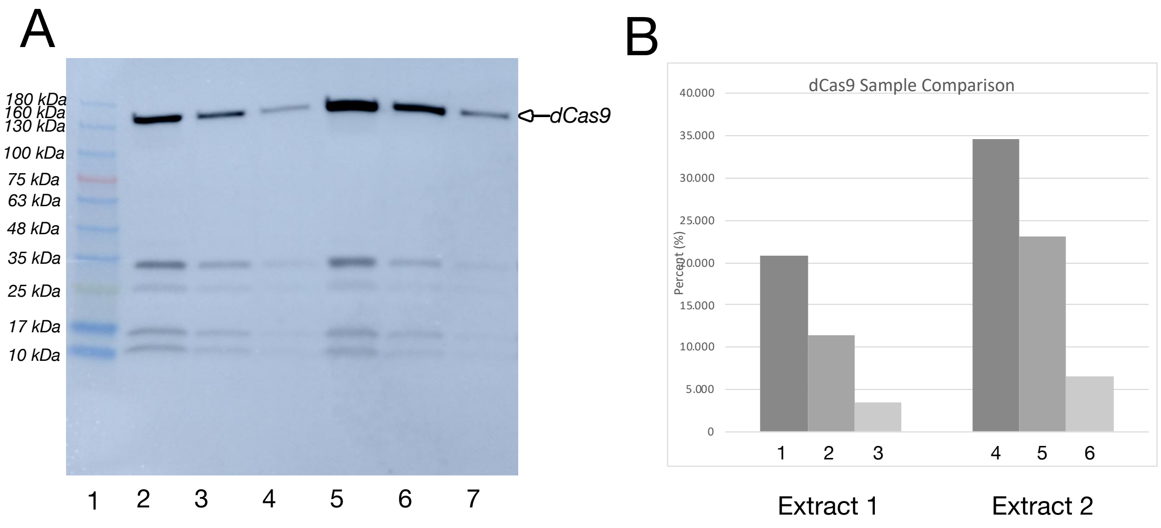


Figure 7: Western blot analysis of two independent batches of dCas9 lysates. Extract 2 corresponds to the dCas9 extract used for quantification in Supp. Fig. 6. A) Line 1: ladder. Line 2: dCas9 lysate-1 at 1:5 dilution. Line 3: dCas9 lysate-1 at 1:10 dilution. Line 4: dCas9-1 lysate at 1:20 dilution. Line 5: dCas9 lysate-2 at 1:5 dilution. Line 6: dCas9 lysate-2 at 1:10 dilution. Line 7: dCas9 lysate-1 at 1:20 dilution. B) Barplot analysis of the reported intensity values analysed in A.

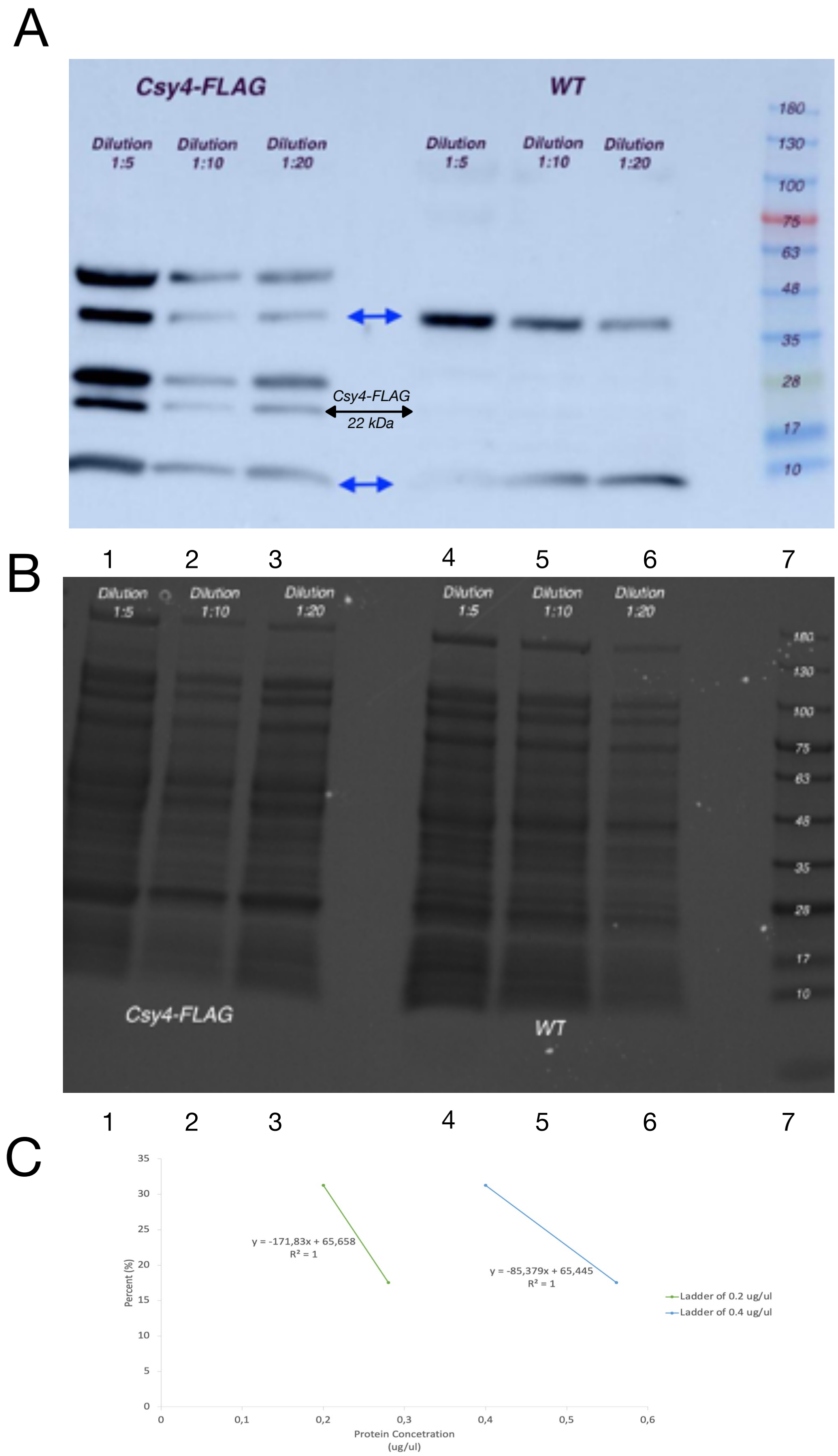


Figure 8: Quantification of Csy4-FLAG in enhanced CFE extract. A) Csy4-FLAG lysate samples (10 µL) were analysed by 4-20% SDS-PAGE, and the separated proteins were transferred onto PVDF membranes probed with anti-Flag antibody. Line 1: Csy4-FLAG lysate at 1:5 dilution. Line 2: Csy4-Flag lysate at 1:10 dilution. Line 3: Csy4-FLAG lysate at 1:5 dilution. Line 4: WT lysate at 1:5 dilution. Line 5: WT lysate at 1:10 dilution. Line 6: WT lysate at 1:20 dilution. Line 7: Ladder B) SDS replicate from A stained by Coomassie Blue G-250. C) Csy4 concentration calculation from ladder (for 0.2 µg and 0.4 µg).

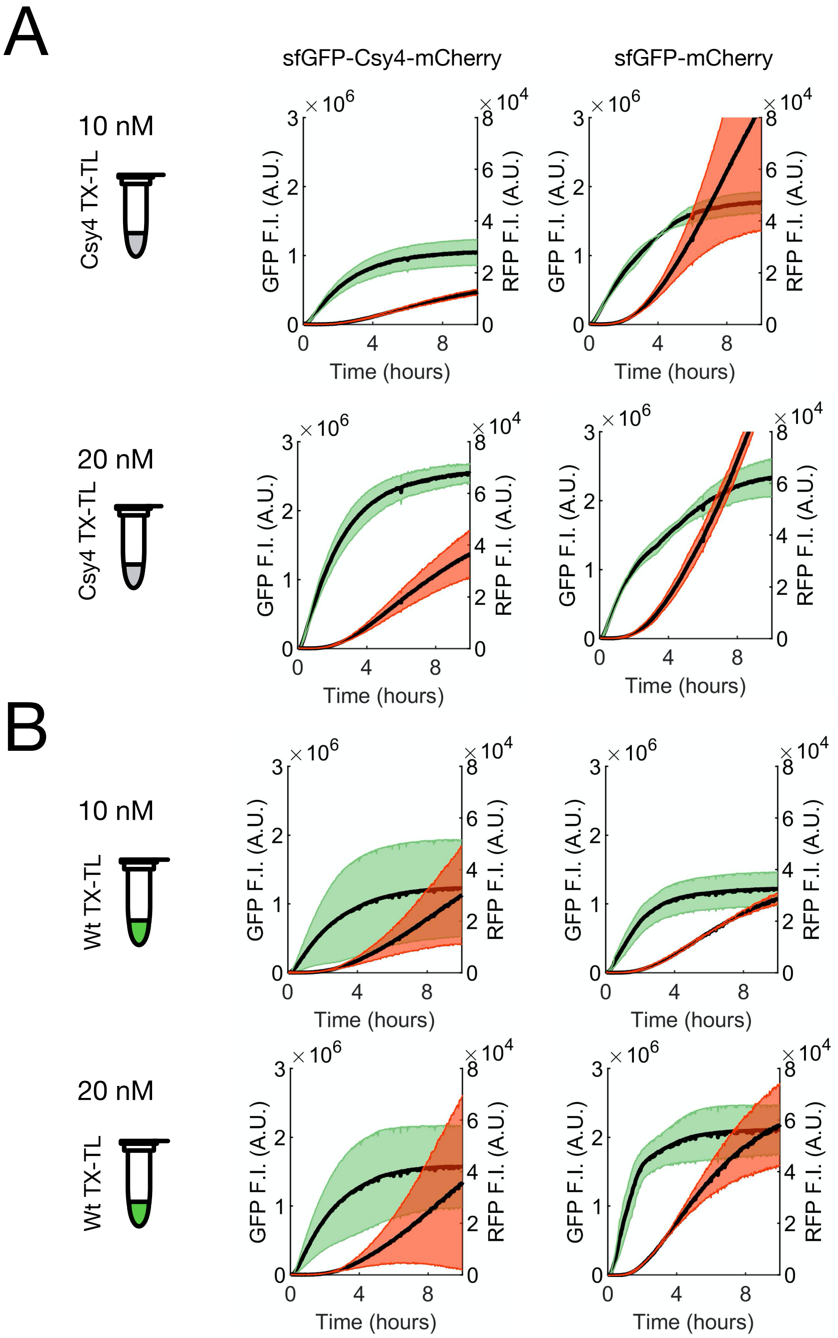


Figure 9: sfGFP (green) and mCherry (red) fluorescent time-courses of sfGFP-Csy4-mCherry and sfGFP-mCherry at 5 nM and 10 nM in Csy4 CFE extract (A) or WT CFE extract (B). Shaded regions represent the standard deviation of the mean from three technical replicates.

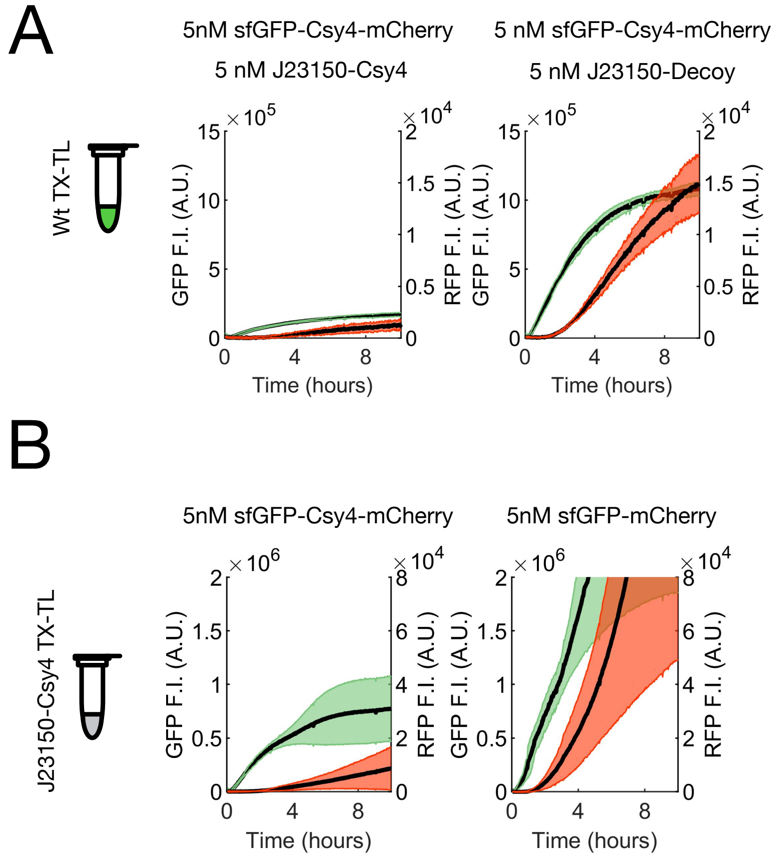


Figure 10: A) sfGFP (green) and mCherry (red) fluorescent time-courses of sfGFP-Csy4-mCherry at 5 nM in WT CFE extract supplemented with 5 nM of Csy4 encoding plasmid or a decoy protein. B) sfGFP (y axis left) and mCherry (y axis right) fluorescent time-courses of sfGFP-Csy4-mCherry and sfGFP-mCherry at 5 nM in J23150-Csy4 CFE extract. Shaded regions represent the standard deviation of the mean from three technical replicates.

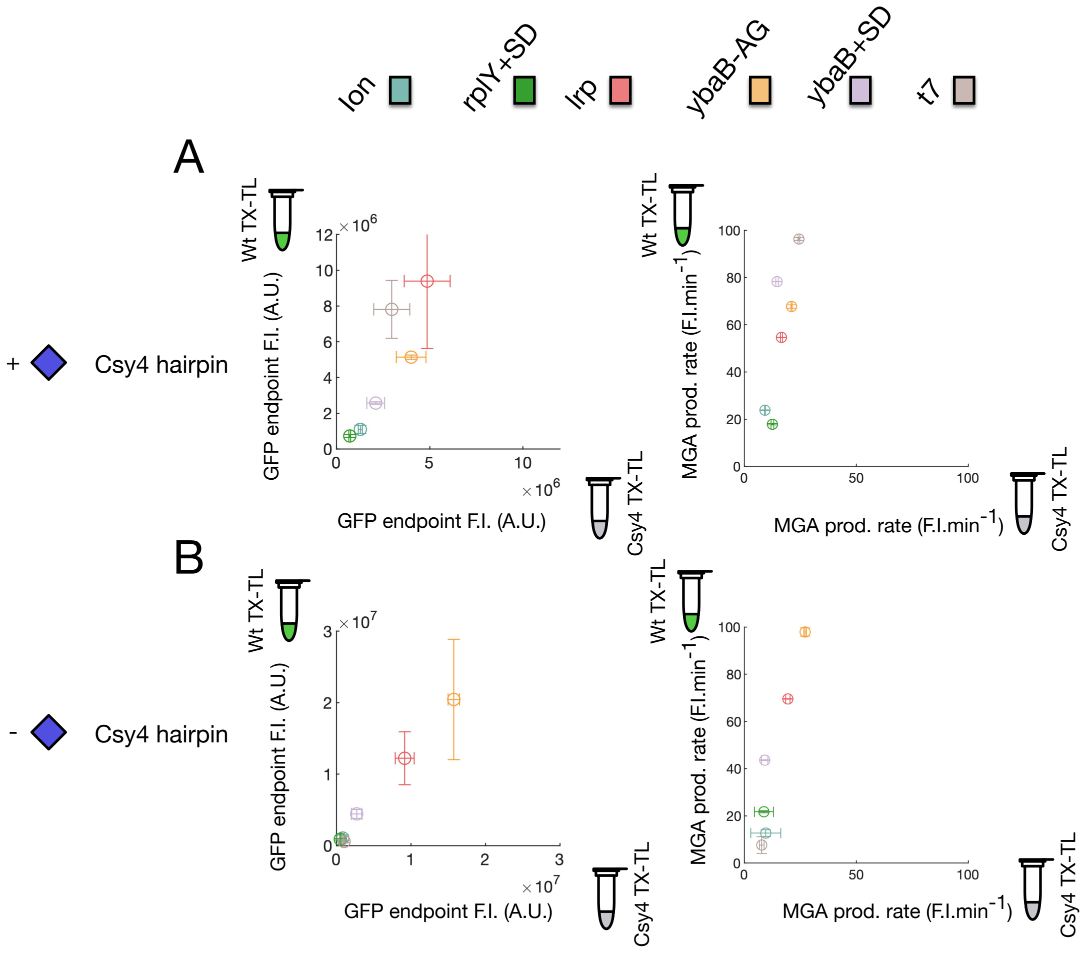


Figure 11: A) sfGFP (left) endpoints and MGA (right) production rates of the six UTR constructs in Csy4 (x axis) or WT (y axis) CFE extract. B) sfGFP (left) endpoints and MGA (right) production rates with Csy4 hairpin constructs in Csy4 (x axis) or WT (y axis) CFE extract. The concentration of UTR constructs was set to 1 nM in all experiments. Error bars and shaded regions represent the standard deviation of the mean from three technical replicates. See Fig. 3 A) for the UTR color legend.

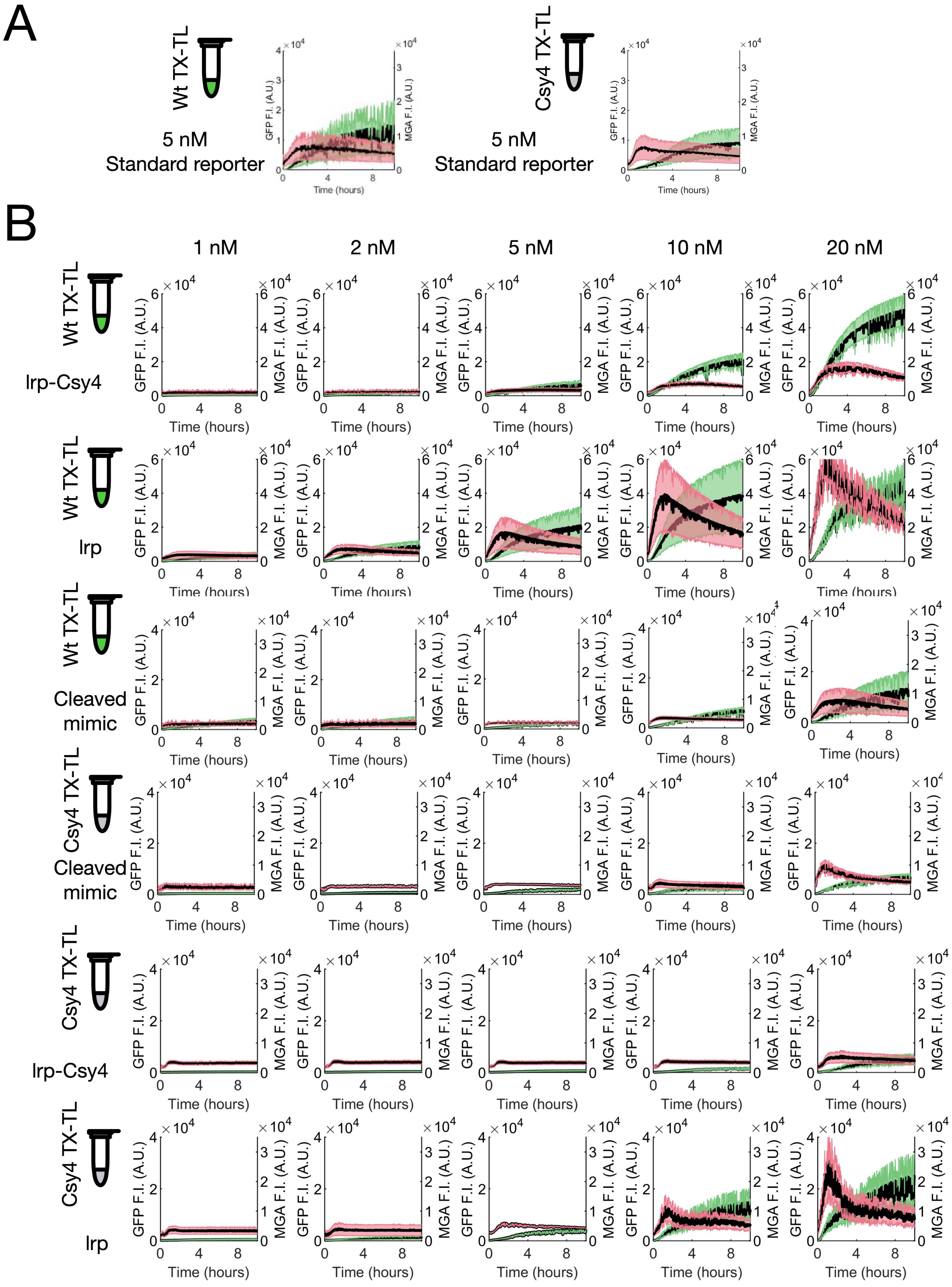


Figure 12: A) sfGFP (green) and MGA (pink) fluorescent time-courses of 5 nM of the standard reporter plasmid in WT and Csy4 extracts. B) sfGFP (green) and MGA (pink) fluorescent time-courses of three constructs (lrp, lrp-Csy4 and the mimic of the cleaved 5' UTR) titrated between 1 and 20 nM in WT and Csy4 CFE extracts. Shaded regions represent the standard deviation of the mean from three technical replicates.

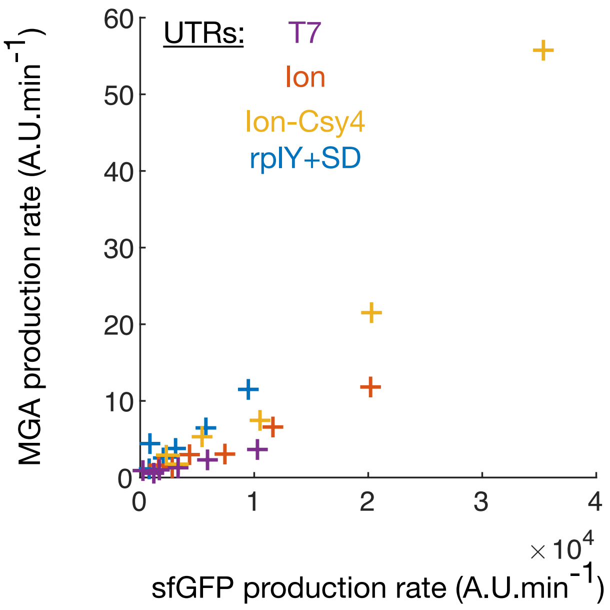


Figure 13: Dose-response relationship of transcription-translation production rates for different UTR constructs. The sfGFP and MGA production rates correspond to the maximum production rate computed over 20 minutes intervals for 10 hours.

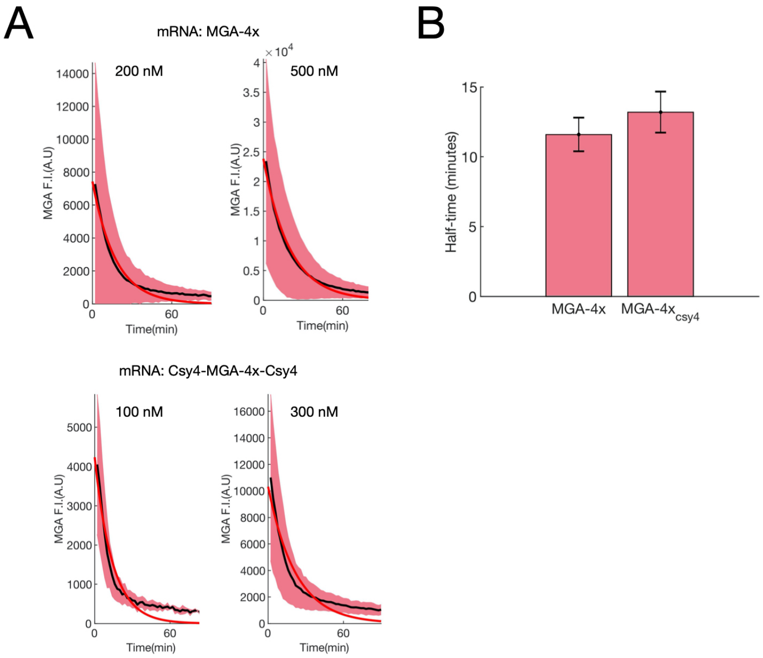


Figure 14: mRNA degradation experiments performed in Csy4 CFE extract. A) Fluorescence decay time traces of purified mRNA titrated (MGA-4x: top, Csy4-MGA-4x-Csy4: bottom) in Csy4 CFE extract for two different concentrations (range = 100-500 nM). Shaded pink regions correspond to the experimental standard variation (n=3). B) Corresponding half-times for two different purified mRNA computed from the decay traces. Purified mRNA standards were prepared in Csy4 CFE extract with 3-PGA energy buffer and 40 µM malachite green. Experimental data-points (black lines) and a non-linear fit (red line) of half-life was estimated based on a single-term exponential model (MatLab 2019a) with $Y = Y_0 e^{-kt}$, where Y is mRNA concentration, $Y_0$ is the mRNA concentration at time zero, $k$ is the decay constant and $t$ is the time constant.

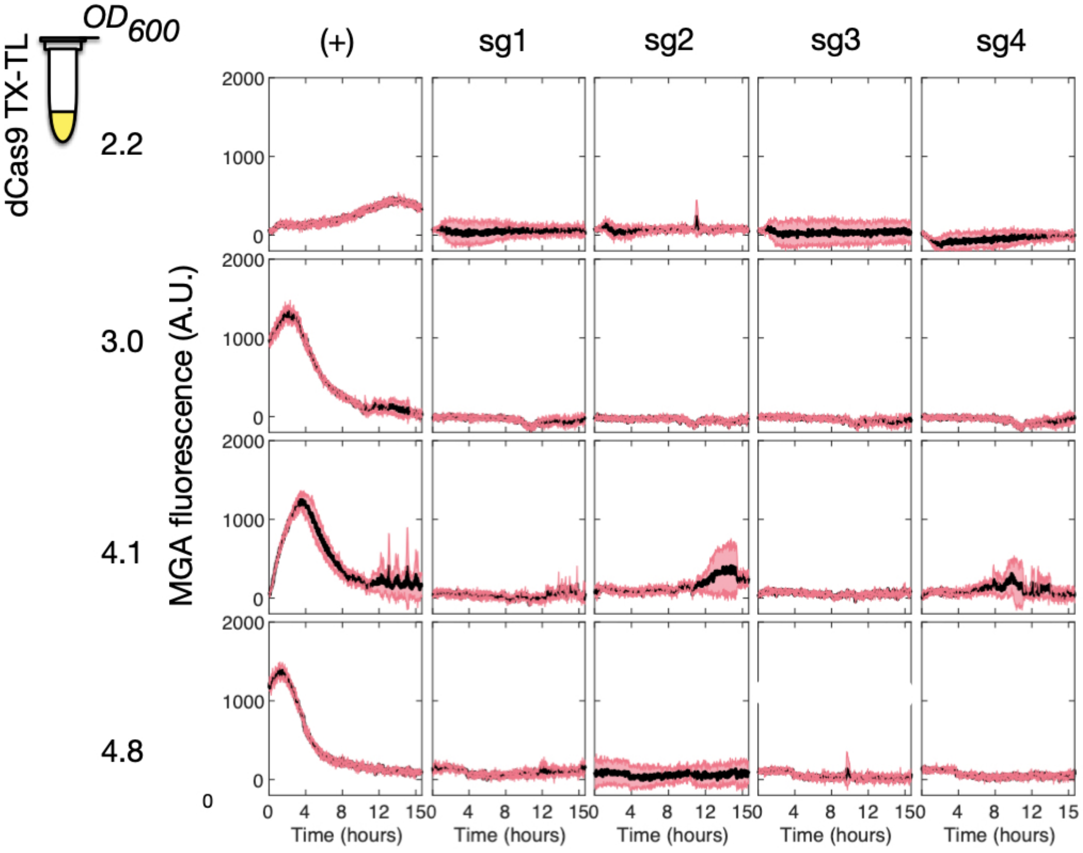


Figure 15: Transcriptional time traces of four dCas9 CFE extracts made from a variety of $OD_{600}$ and in presence of 10 nM of each sgRNA. The concentration of reporter plasmid (Pr-sfGFP) was set to 1 nM in all experiments. The red shaded regions correspond to the standard deviation over three independent reactions.

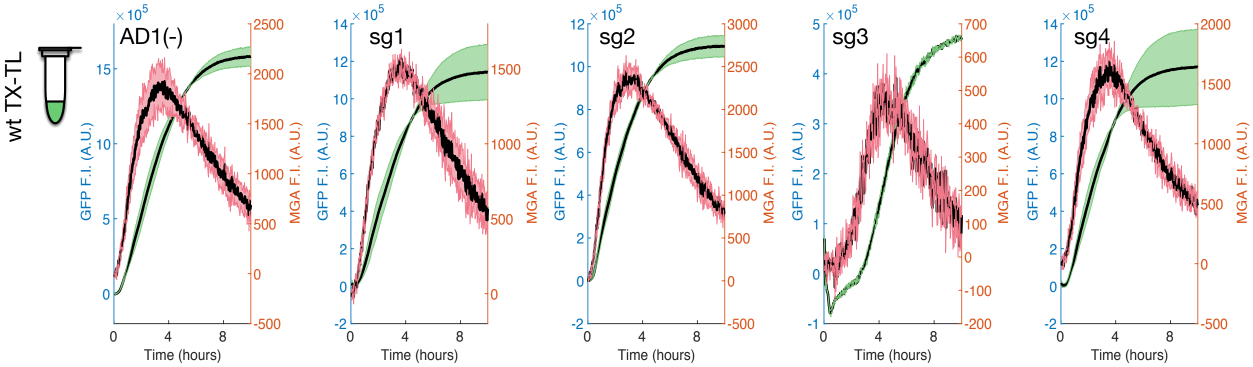


Figure 16: Expression of the sgRNAs does not inhibit gene expression in wild-type CFE extract. Fluorescent MGA and sfGFP time traces of 1 nM of Pr-sfGFP plasmid in presence of 10 nM of AD1/Sg-1/Sg-2/Sg-3/Sg-4 plasmid. AD1 corresponds to a decoy sRNA used to control if the transcriptional burden of the sRNA inhibits the sGFP production. The green and red shaded regions correspond to the standard deviation over three independent reactions.

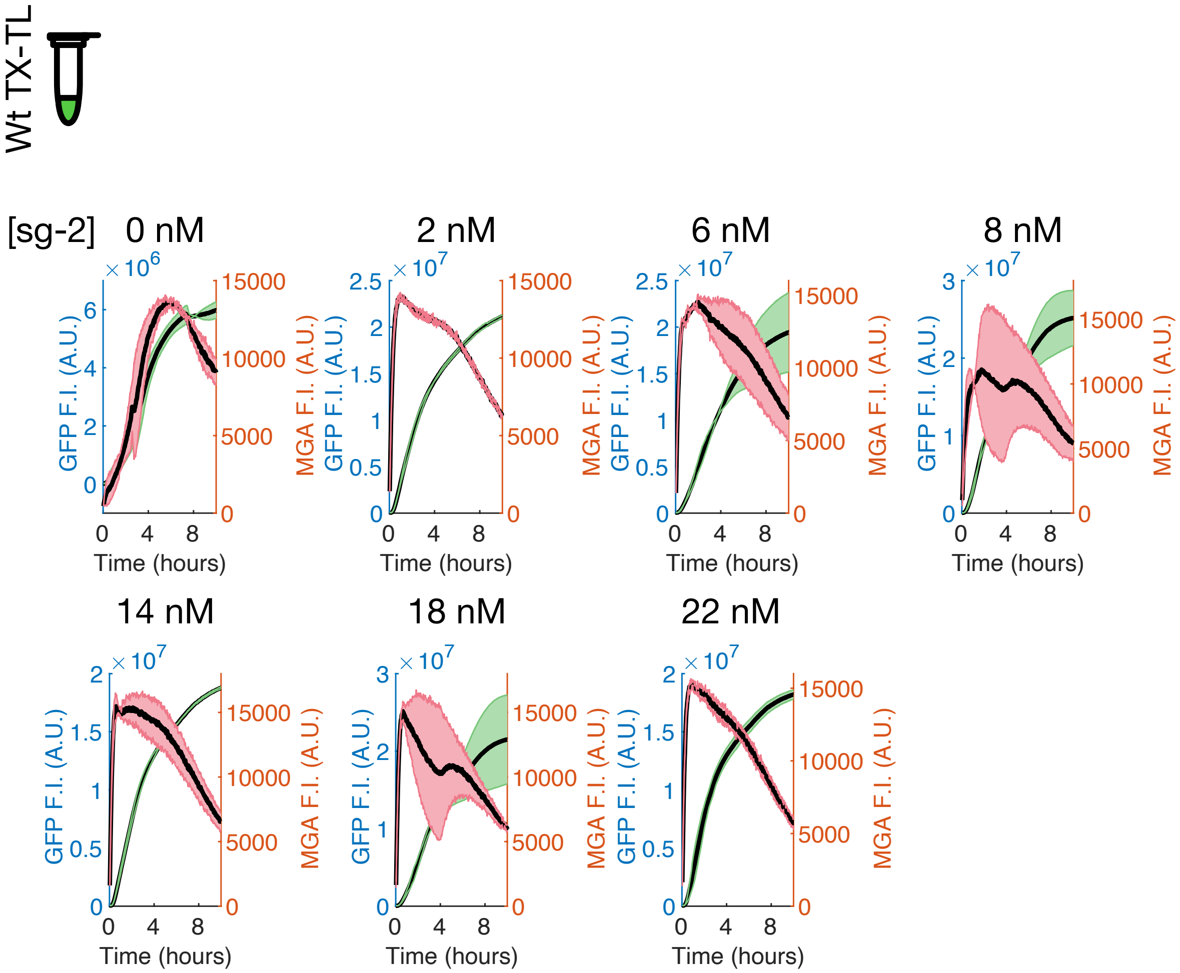


Figure 17: Expression of the sgRNAs does not inhibit gene expression in wild-type CFE extract. Fluorescent MGA and sfGFP time traces of 8 nM of Pr-sfGFP plasmid titrated with a range from 0 nM to 22 nM of Sg-2 plasmid

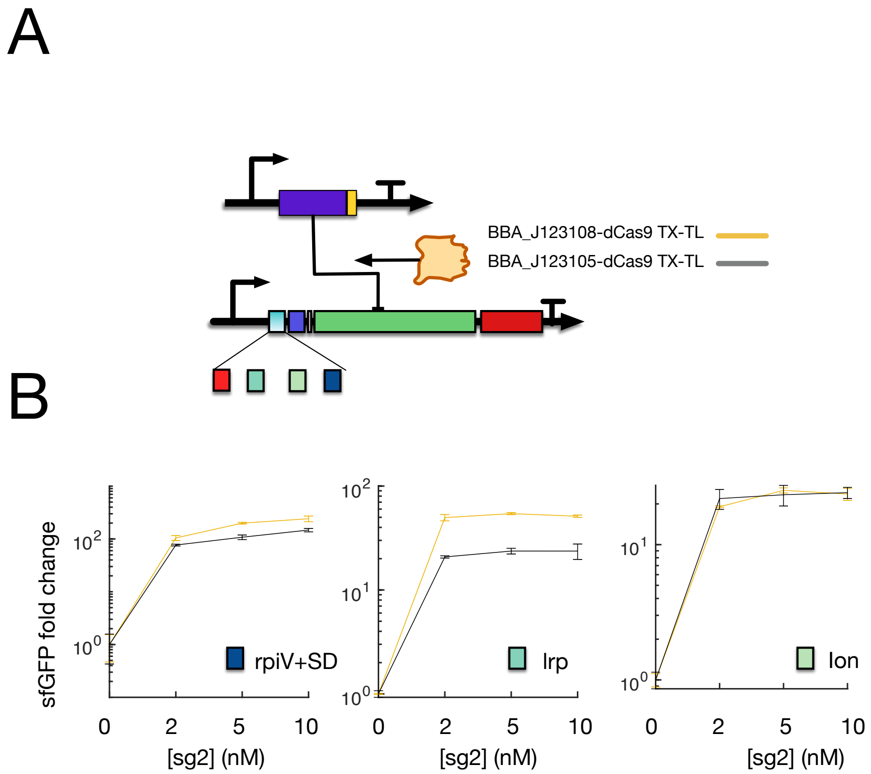


Figure 18: A) Scheme of the dCas9 regulation mechanism. B) sfGFP fold-change repressions for three UTR constructs titrated with sgRNA 2 plasmid, performed in dCas9 CFE extracts expressed from two different promoters. Error bars represent the standard deviation of the mean from three technical replicates

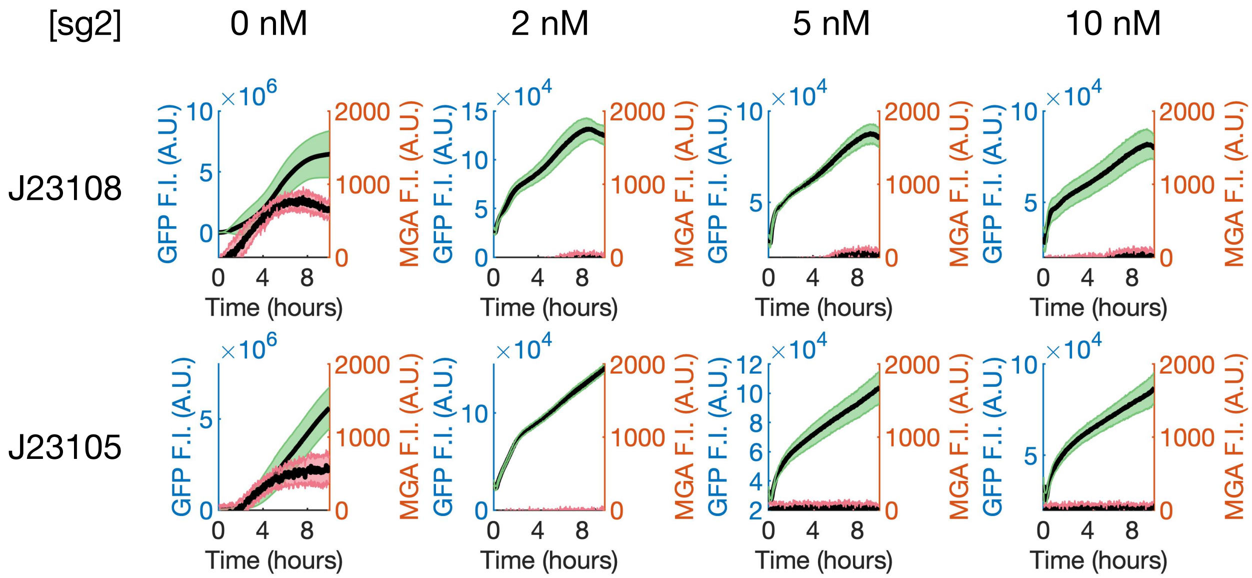


Figure 19: Fluorescent MGA and sfGFP traces for 1 nM of Ion-UTR construct in presence of a range of 0-10 nM of sg-2 plasmid. The top and bottom row display kinetic experiments performed in J23108-dCas9 J23105-dCas9 CFE extracts, respectively. The green and red shaded regions correspond to the standard deviation over three independent reactions.

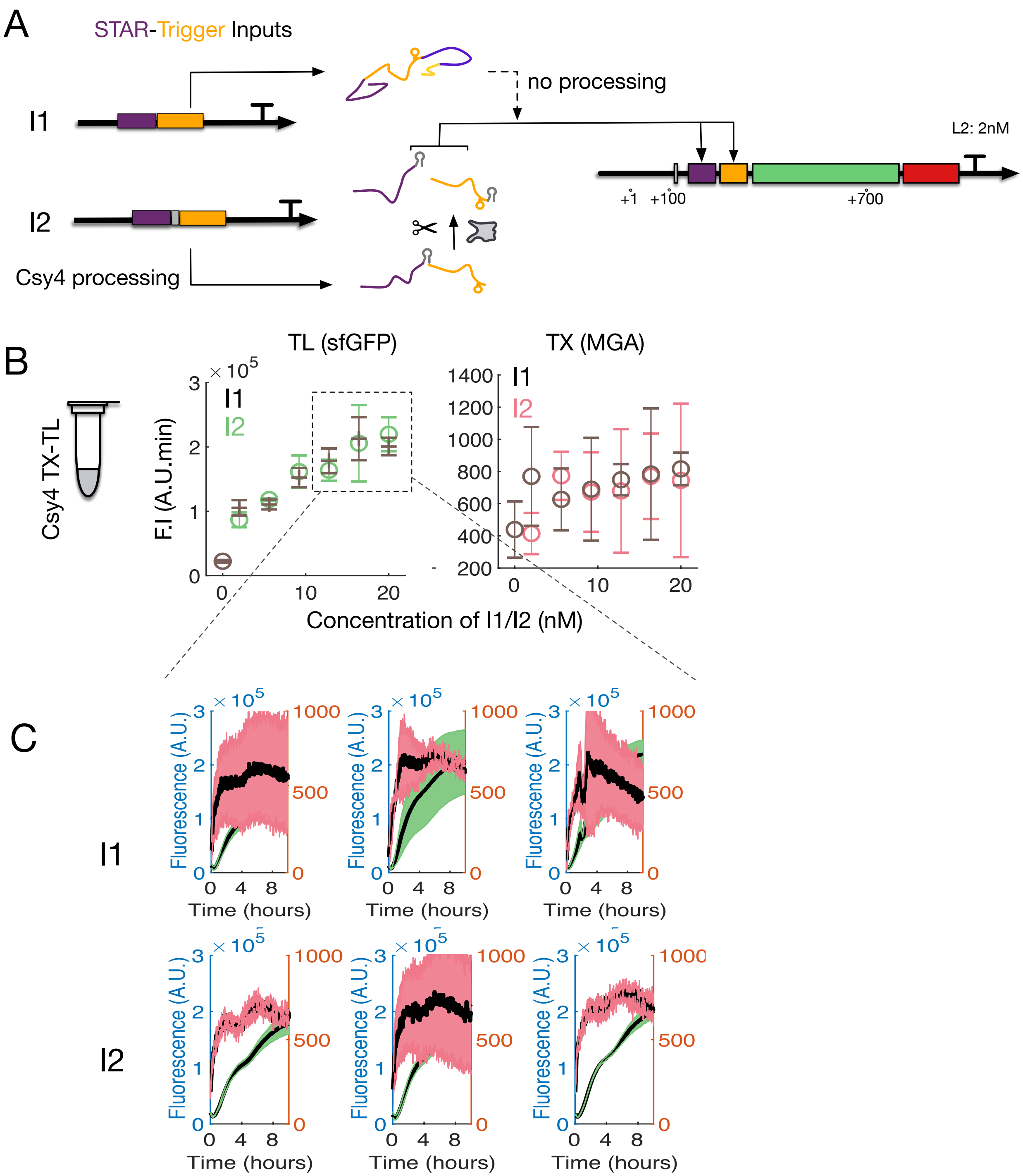


Figure 20: Trans-activation of the S6T3 gate is not improved by Csy4 processing. A) Scheme of the trans-acting sRNA operon constructs, I-1 and I-2. B) Endpoint (sfGFP) and maximum (MGA) fluorescent measurements for I-1 and I-2 in presence of 2 nM of S6T3 gate reporter. C) Corresponding fluorescence trace examples for three concentrations from B). Errors bars represent the standard deviation over three replicates.

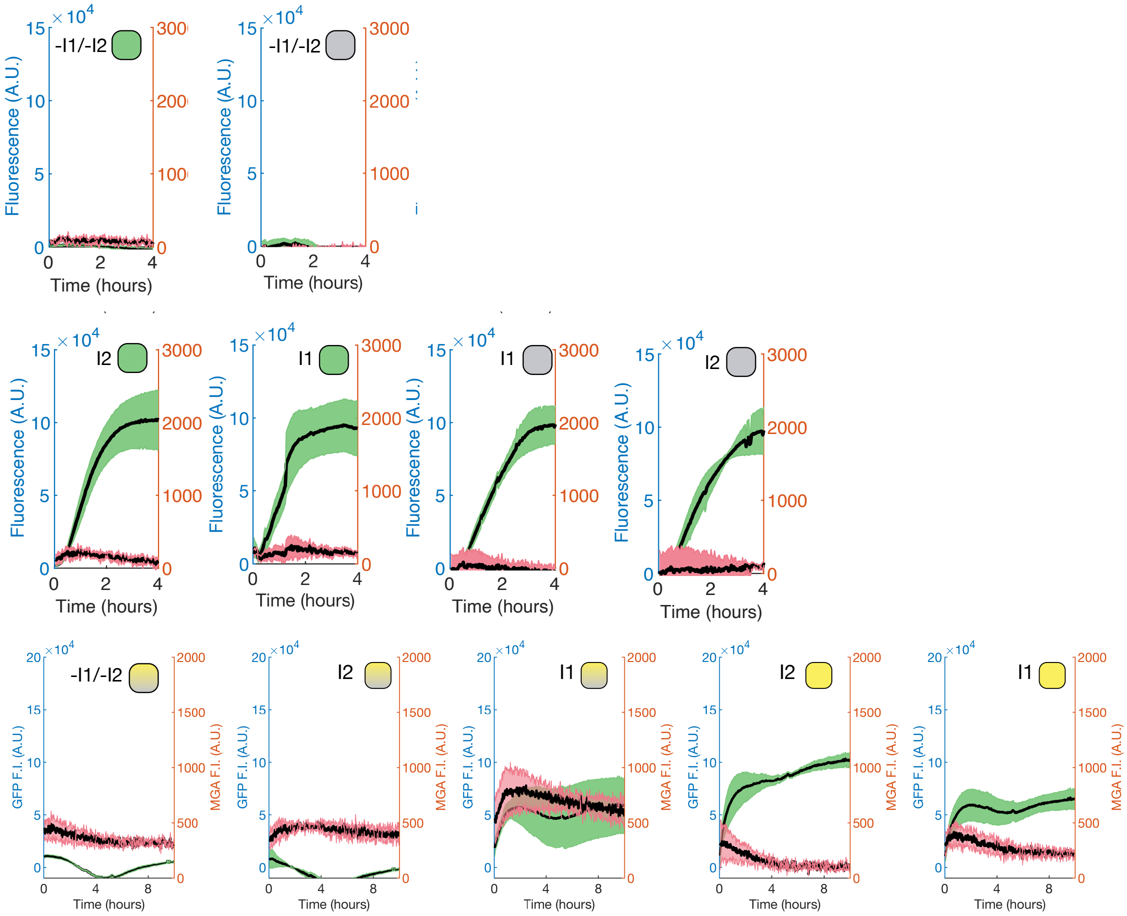


Figure 21: sfGFP and MGA fluorescent time traces used for the sRNA-operon experiment with 2 nM of S6T3 gate plasmid (-I1/-I2, orange) and 10 nM of either I1 (grey) or I2 (blue) in four different CFE extracts. Csy4 (grey) and dCas9 (yellow) CFE extracts are mixed to form a ”blended” CFE extract (shaded). Shaded regions represent the standard deviation of the mean from three technical replicates.

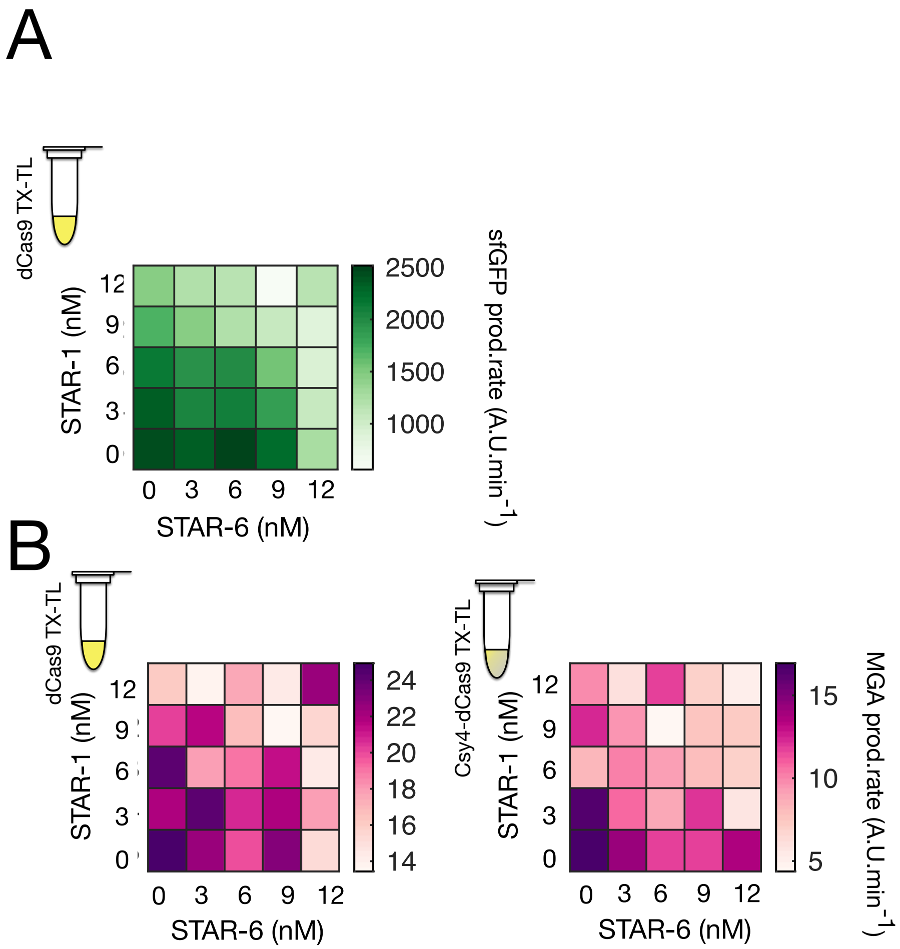


Figure 22: A) Heatmap of the sfGFP production rates for the NAND gate titrated with STAR-1 and STAR-6 encoding plasmids in dCas9 extract. B) Heatmaps of the MGA production rates for the NAND gate titrated with STAR-1 and STAR-6 encoding plasmids in either dCas9 (left) or blended Csy4-dCas9 (right) extract. Other parts concentrations were set to 2 nM (reporter plasmid) and 10 nM (sg-2 expressing plasmid).

Table 1: Plasmids used.

| Plasmid Name | Plasmid architecture | Source |
|---|---|---|
| pJBL2801 | J23119 – Target AD1.S5 – RBS – sfGFP – TrrnB – CmR – p15A origin | Addgene 71207 |
| pJBL-wt | J23119 — RBS – sfGFP – TrrnB – CmR – p15A origin | this study |
| pJBL-Toehold2 | J23119 – Toehold2 – RBS – sfGFP – TrrnB – CmR – p15A origin | this study |
| pJBL-trigger2 | J23119 – trigger2 – T500 – AmpR – ColE1 origin | this study |
| pJBL-trigger3 | J23119 – trigger3 – T500 – AmpR – ColE1 origin | this study |
| pJBL-Target6-Toehold2 | J23119 – Target6 – Toehold2 – sfGFP – TrrnB – CmR – p15A origin | this study |
| pJBL-Target6-Toehold3 | J23119 – Target6 – Toehold3 – sfGFP – TrrnB – CmR – p15A origin | this study |
| pJBL-Target6-Target1-sgRNA2-C | J23119 – Target6 – Csy4 – Target1 – Csy4 – sgRNA2 – T500 – AmpR – ColE1 origin | this study |
| pJBL-STAR AD1.A5-trigger2 | t500 – trigger2 – J23119 – spacer – J23119 – STAR AD1.A5 – T500 – AmpR – ColE1 origin | this study |
| pJBL-STAR6-trigger2 | t500 – trigger2 – J23119 – spacer – J23119 – STAR6 – T500 – AmpR – ColE1 origin | this study |
| pJBL-STAR6-trigger3 | t500 – trigger3 – J23119 – spacer – J23119 – STAR6 – T500 – AmpR – ColE1 origin | this study |
| pFXL-S6T3-MGA | J23119 – Target6 – Toehold3 – sfGFP-MGA4x – T500 – Amp – ColE1 origin | this study |
| pFXL-S6T2-MGA | J23119 – Target6 – Toehold2 – sfGFP-MGA4x – T500 – Amp – ColE1 origin | this study |
| pFXL-S6T2C-MGA | J23119 – Target6 – Csy4 – Toehold2 – sfGFP-MGA4x – T500 – Amp – ColE1 origin | this study |
| pFXL-108-dCas9 | J23108 – RBS – SpydCas9 – T500 – AmpR – ColE1 origin | this study |
| pFXL-107-dCas9 | J23107 – RBS – SpydCas9 – T500 – AmpR – ColE1 origin | this study |
| pFXL-115-dCas9 | J23115 – RBS – SpydCas9 – T500 – AmpR – ColE1 origin | this study |
| pFXL-107-Csy4 | J23107 – RBS – Csy4 – T500 – AmpR – ColE1 origin | this study |
| pFXL-115-Csy4 | J23115 – RBS – Csy4 – T500 – AmpR – ColE1 origin | this study |
| pFXL-108-Csy4 | J23108 – RBS – Csy4 – T500 – AmpR – ColE1 origin | this study |
| pFXL-deCas9-Csy4 | J23108 – RBS – Csy4 – T500 – J23108 – RBS – deCas9 – T500 – AmpR – ColE1 origin | this study |
| pJBL-sgRNA1 | J23119 – sgRNA1 – T500 – AmpR – ColE1 origin | this study |
| pJBL-sgRNA2 | J23119 – sgRNA2 – T500 – AmpR – ColE1 origin | this study |
| pJBL-sgRNA3 | J23119 – sgRNA3 – T500 – AmpR – ColE1 origin | this study |
| pJBL-sgRNA4 | J23119 – sgRNA4 – T500 – AmpR – ColE1 origin | this study |

| Plasmid Name | Plasmid architecture | Source |
|---|---|---|
| pJBL-sg2-s6T3-C | J23119 – STAR6 – Cys4 - Trigger3 – Cys4 – sgRNA2 – T500 – AmpR – ColE1 origin | this study |
| pJBL-sg2-s6T3 | J23119 – STAR6 – Trigger3 – sgRNA2 – T500 – AmpR – ColE1 origin | this study |
| pFXL02-ion | J23119 – ion– RBS – sfGFP-MGA4x – T500 – amp – ColE1 origin | this study |
| pFXL02-ionC- | J23119 – ion– Csy4 – RBS – sfGFP-MGA4x – T500 – amp – ColE1 origin | this study |
| pFXL02-rpiY-SD | J23119 – rpiY-SD– RBS – sfGFP-MGA4x – T500 – amp – ColE1 origin | this study |
| pFXL02-rpiY-SD-C | J23119 – rpiY-SD– Csy4 – RBS – sfGFP-MGA4x – T500 – amp – ColE1 origin | this study |
| pFXL02-t7 | J23119 – t7– RBS – sfGFP-MGA4x – T500 – amp – ColE1 origin | this study |
| pFXL02-t7C- | J23119 – t7– Csy4 – RBS – sfGFP-MGA4x – T500 – amp – ColE1 origin | this study |
| pFXL02-ybaB-AG | J23119 – ybaB-AG– RBS – sfGFP-MGA4x – T500 – amp – ColE1 origin | this study |
| pFXL02-ybaB-AG-C | J23119 – ybaB-AG– Csy4 – RBS – sfGFP-MGA4x – T500 – amp – ColE1 origin | this study |
| pFXL02-Irp | J23119 – Irp– RBS – sfGFP-MGA4x – T500 – amp – ColE1 origin | this study |
| pFXL02-Irp-C | J23119 – Irp– Csy4 – RBS – sfGFP-MGA4x – T500 – amp – ColE1 origin | this study |
| pFXL02-ybaB-SD | J23119 – ybaB-SD– RBS – sfGFP-MGA4x – T500 – amp – ColE1 origin | this study |
| pFXL02-ybaB-SD-C | J23119 – ybaB-SD– Csy4 – RBS – sfGFP-MGA4x – T500 – amp – ColE1 origin | this study |
| pFXL02-cleaved-mimc | J23119 – partial csy4 hairpin – RBS – sfGFP – RBS – mCherry – T7 terminator – amp – pBR322 origin | this study |
| GFP-RFP | J23119 – RBS – sfGFP – RBS – mCherry – T7 terminator – amp – pBR322 origin | this study |
| GFP-Csy4-RFP | J23119 – RBS – sfGFP – RBS – Csy4 – mCherry – T7 terminator – amp – pBR322 origin | this study |

Table 2: Example Plasmids.

| Name | Sequence [5'->3'] |
|---|---|
| pJBL-STAR5-trigger1 t500, trigger, J23119, STAR, ColE1, AmpR | GAATTCaaaaaaaagcccgcctttcggcgggctttgAGATCCctatcttatcttatctatctcgtttatccctgctttactgact<br>attgcacagaatagtcagtcccactagtattatacctaggactgagctagctgtcaaAGATCTTTAACGGGGTCAT<br>CACGGCTCATCATGCGCCAAACAAATGTGTGCAATACACGCTCGGATGACTGCA<br>TGATGACCGCACTGACTGGGGACAGCAGATCCACCTAAGCCTGTGAGAGAAGCA<br>GACACCCGACAGATCAAGGCAGTTAAATTAAAGATCTttgacagctagctcagtcctaggtat<br>aatactagttgaactgtatacattccccgcaggataggaattgaagatgaaacgatgagacttgggacgaGGATCT<br>caaagcccgccgaaaggcgggcttttttttGGATCCTTACTCGAGTCTAGACTGCAGGCTTCCT<br>CGCTCACTGACTCGCTGCGCTCGGTCGTTCGGCTGCGGCGAGCGGTATCAGCTC<br>ACTCAAAGGCGGTAATACGGTTATCCACAGAATCAGGGGATAACGCAGGAAAGA<br>ACATGTGAGCAAAAGGCCAGCAAAAGGCCAGGAACCGTAAAAAggccgcgttgctggcg<br>tttttccacaggctccgcccccctgacgagcatcacaaaaatcgacgctcaagtcagaggtggcgaaacccgacaggactataaa<br>gataccaggcgtttccccctggaagctccctcgtgcgctctcctgttccgaccctgccgcttaccggatacctgtccgcctttct<br>cccttcgggaagcgtggcgctttctcatagctcacgctgtaggtatctcagttcggtgtaggtcgttcgctccaagctgggctgt<br>gtgcacgaaccccccgttcagcccgaccgctgcgccttatccggtaactatcgtcttgagtccaacccggtaagacacgacttat<br>cgccactggcagcagccactggtaacaggattagcagagcgaggtatgtaggcggtgctacagagttcttgaagtggtggcctaa<br>ctacggctacactagaagaacagtatttggtatctgcgctctgctgaagccagttaccttcggaaaaagagttggtagctcttga<br>tccggcaaacaaaccaccgctggtagcggtggtttttttgtttgcaagcagcagattacgcgcagaaaaaaaggatctcaagaag<br>atcctttgatcttttctacggggtctgacgctcagtggaacgaaaactcacgttaagggattttggtcatgaGATTATCA<br>AAAAGGATCTTCACCTAGATCCTTTTAAATTAAAAATGAAGTTTTAAATCATCT<br>AAAGTATATATGAGTAAACTTGGTCTGACAGTTAccaatgcttaatcagtgaggcacctatctcagc<br>gatctgtctatttcgttcatccatagttgcctgactccccgtcgtgtagataactacgatacgggagggcttaccatctggcccc<br>agtgctgcaatgataccgcgagacccacgctcaccggctccagatttatcagcaataaaccagccagccggaagggccgagcgca<br>gaagtggtcctgcaactttatccgcctccatccagtctattaattgttgccgggaagctagagtaagtagttcgccagttaatag |
| pJBL-STAR5-trigger1 t500, trigger, J23119, STAR, ColE1, AmpR | tttgcgcaacgttgttgccattgctacaggcatcgtggtgtcacgctcgtcgtttggtatggcttcattcagctccggttcccaa<br>cgatcaaggcgagttacatgatcccccatgttgtgcaaaaaagcggttagctccttcggtcctccgatcgttgtcagaagtaagt<br>tggccgcagtgttatcactcatggttatggcagcactgcataattctcttactgtcatgccatccgtaagatgcttttctgtgac<br>tggtgagtactcaaccaagtcattctgagaatagtgtatgcggcgaccgagttgctcttgcccggcgtcaatacgggataatacc<br>gcgccacatagcagaactttaaaagtgctcatCATTGGAAAACGTTCTTCGGGGCGAAAACTCTCA<br>AGGATCTTACCGCTGTTGAGATCCAGTTCGATGTAACCCACTCGTGCACCCAACTG<br>ATCTTCAGCATCTTTTACTTTCACCAGCGTTTCTGGGTGAGCAAAAACAGGAAGGC<br>AAAATGCCGCAAAAAAGGGAATAAGGGCGACACGGAAATGTTGAATACTCATACTC<br>TTCCTTTTTCAATATTATTGAAGCATTTATCAGGGTTATTGTCTCATGAGCGGATA<br>CATATTTGAATGTATTTAGAAAAATAAACAAATAGGGGTTCCGCGCACATTTCCCC<br>GAAAAGTGCCACCTGACGTCTAAGAAACCATTATTATCATGACATTAACCTATAAA<br>AATAGGCGTATCACGAGGCAGAATTTCAGATAAAAAAAATCCTTAGCTTTCGCTAA<br>GGATGATTTCTG |

| **Name** | **Sequence [5’->3’]** |
|---|---|
| pJBL-Target5-<br>Toehold1<br>J23119,<br>Target,<br>toehold,<br>sfGFP, TrrnB,<br>CmR, p15A | GAATTCTAAAGATCTttgacagctagctcagtcctaggtataatactagttcgtcccaagtctcatcgtttcatct<br>tcaattcctatcctgcggggaatgtatacagttcatgtatatattccccgctttttttttGGATCTgggtcttatcttatc<br>tatctcgtttatccctgcatacagaaacagaggagatatgcaatgataaacgagaacctggcggcagcgcaaaag<br>atgagcaaaggagaagaacttttcactggagttgtcccaattcttgttgaattagatggtgatgttaatgggcacaaattttctgtc<br>cgtggagagggtgaaggtgatgctacaaacggaaaactcacccttaaatttatttgcactactggaaaactacctgttccgtgg<br>ccaacacttgtcactactctgacctatggtgttcaatgcttttcccgttatccggatcacatgaaacggcatgactttttcaag<br>agtgccatgcccgaaggttatgtacaggaacgcactatatctttcaaagatgacgggacctacaagacgcgtgctgaagtcaag<br>tttgaaggtgatacccttgttaatcgtatcgagttaaagggtattgattttaaagaagatggaaacattcttggacacaaactc<br>gagtacaactttaactcacacaatgtatacatcacggcagacaaacaaaagaatggaatcaaagctaacttcaaaattcgccac<br>aacgttgaagatggttccgttcaactagcagaccattatcaacaaaatactccaattggcgatggccctgtccttttaccagac<br>aaccattacctgtcgacacaatctgtcctttcgaaagatcccaacgaaaagcgtgaccacatggtccttcttgagtttgtaact<br>gctgctgggattacacatggcatggatgagctctacaaaTAAGCGGCCGCGGATCTgaagcttgggcccgaaca<br>aaaactcatctcagaagaggatctgaatagcgccgtcgaccatcatcatcatcatcattgagtttaaacggtctccagcttggc<br>tgttttggcggatgagagaagattttcagcctgatacagattaaatcagaacgcagaagcggtctgataaaacagaatttgcct<br>ggcggcagtagcgcggtggtcccacctgaccccatgccgaactcagaagtgaaacgccgtagcgccgatggtagtgtggggtct<br>ccccatgcgagagtagggaactgccaggcatcaaataaaacgaaaggctcagtcgaaagactgggcctttcgttttatctgttg<br>tttgtcggtgaactGGATCCTTACTCGAGTCTAGACTGCAGTTGATCGggcacgtaagaggttc<br>caactttcaccataatgaaataagatcactaccgggcgtattttttgagttatcgagattttcaggagctaaggaagctaaaat<br>ggagaaaaaaatcactggatataccaccgttgatatatcccaatggcatcgtaaagaacattttgaggcatttcagtcagttgc<br>tcaatgtacctataaccagaccgttcagctggatattacggcctttttaaagaccgtaaagaaaaataagcacaagttttatcc<br>ggcctttattcacattcttgcccgcctgatgaatgctcatccggaatttcgtatggcaatgaaagacggtgagctggtgatatg<br>ggatagtgttcacccttgttacaccgttttccatgagcaaactgaaacgttttcatcgctctggagtgaataccacgacgattt<br>ccggcagtttctacacatatattcgcaagatgtggcgtgttacggtgaaaacctggcctatttccctaaagggtttattgagaa<br>tatgtttttcgtctcagccaatccctgggtgagtttcaccagttttgatttaaacgtggccaatatggacaacttcttcgcccc<br>cgttttcaccatgggcaaatattatacgcaaggcgacaaggtgctgatgccgctggcgattcaggttcatcatgccgtttgtga<br>tggcttccatgtcggcagaatgcttaatgaattacaacagtactgcgatgagtggcagggcggggcgtaatttgatatcgagct<br>cgcttggactcctgttgatagatccagtaatgacctcagaactccatctggatttgttcagaacgctcggttgccgccgggcgt<br>tttttattGGTGAGAATCCAAGCCTCCGATCAACGTCTCATTTTCGCCAAAAGTTG |

| **Name** | **Sequence [5'->3']** |
|---|---|
| pJBL-Target5-Toehold1 J23119, Target, toehold, sfGFP, TrrnB, CmR, p15A | GCCCAGGGCTTCCCGGTATCAACAGGGACACCAGGATTTATTTATTCTGCGAAGT GATCTTCCGTCACAGGTATTTATTCGGCGCAAAGTGCGTCGGGTGATGCTGCCAA CTTACTGATTTAGTGTATGATGGTGTTTTTGAGGTGCTCCAGTGGCTTCTGTTTC TATCAGCTGTCCCTCCTGTTCAGCTACTGACGGGGTGGTGCGTAACGGCAAAAGC ACCGCCGGACATCAgcgctagcggagtgtatactggcttactatgttggcactgatgagggtgtcagtgaagtgcttc atgtggcaggagaaaaaaggctgcaccggtgcgtcagcagaatatgtgatacaggatatattccgcttcctcgctcactgactc gctacgctcggtcgttcgactgcggcgagcggaaatggcttacgaacggggcggagatttcctggaagatgccaggaagatact taacagggaagtgagagggccgcggcaaagccgtttttccataggctccgcccccctgacaagcatcacgaaatctgacgctca aatcagtggtggcgaaacccgacaggactataaagataccaggcgtttccccctggcggctccctcgtgcgctctcctgttcct gcctttcggtttaccggtgtcattccgctgttatggccgcgtttgtctcattccacgcctgacactcagttccgggtaggcagt tcgctccaagctggactgtatgcacgaaccccccgttcagtccgaccgctgcgccttatccggtaactatcgtcttgagtccaa cccggaaagacatgcaaaagcaccactggcagcagccactggtaattgatttagaggagttagtcttgaagtcatgcgccggtt aaggctaaactgaaaggacaagttttggtgactgcgctcctccaagccagttacctcggttcaaagagttggtagctcagagaa ccttcgaaaaaccgccctgcaaggcggttttttcgttttcagagcaagagattacgcgcagaccaaaacgatctcaagaagatc atcttattaatcagataaaatatttCTAGATTTCAGTGCAATTTATCTCTTCAAATGTAGCACC TGAAGTCAGCCCCATACGATATAAGTTGTAATTCTCATGTTTGACAGCTTATCAT CGATAAGCTTCCGATGGCGCGCCGAGAGGCTTTACACTTTATGCTTCCGGCT |

| Name | Sequence [5'->3'] |
|---|---|
| sfGFP-Csy4-mCherry J23119, RBS, sfGFP, Csy4, RBS, mCherry, , T500 | TTGACAGCTAGCTCAGTCCTAGGTATAATACTAGTATGTCTTCGGATCTTAGCTACTAG<br>AGAAAGAGGAGAAATACTAGATGCGTAAAGGCGAAGAGCTGTTCACTGGTGTCGTCCC<br>TATTCTGGTGGAACTGGATGGTGATGTCAACGGTCATAAGTTTTCCGTGCGTGGC-<br>GAGGGTGAA<br>GGTGACGCAACTAATGGTAAACTGACGCTGAAGTTCATCTGTACTACTGGTAAAC<br>TGCCGGTACCTTGGCCGACTCTGGTAACGACGCTGACTTATGGTGTTCAGTGCTTT<br>GCTCGTTATCCGGACCATATGAAGCAGCATGACTTCTTCAAGTCCGCCATGCCGGAA<br>GGCTATGTGCAGGAACGCACGATTTCCTTTAAGGATGACGGCACGTACAAAACGCG<br>TGCGGAAGTGAAATTTGAAGGCGATACCCTGGTAAACCGCATTGAGCTGAAAGGC<br>ATTGACTTTAAAGAAGACGGCAATATCCTGGGCCATAAGCTGGAATACAATTTTAAC<br>AGCCACAATGTTTACATCACCGCCGATAAACAAAAAAATGGCATTAAAGCGAATT<br>TTAAAATTCGCCACAACGTGGAGGATGGCAGCGTGCAGCTGGCTGATCACTACCA<br>GCAAAACACTCCAATCGGTGATGGTCCTGTTCTGCTGCCAGACAATCACTATCTGAG<br>CACGCAAAGCGTTCTGTCTAAAGATCCGAACGAGAAACGCGATCATATGGTTCTGCT<br>GGAGTTCGTAACCGCAGCGGGCATCACGCATGGTATGGATGAACTGTACAAATAAG<br>CGGATCCGAATAATTTTGTTTAACTTTAAGAAGGAGATATACATATGCGTGAGTAAAGG<br>CGAGGAGGACAATATGGTTCACTGCCGTATAGGCAGGCGATCATCAAAGAGTTC<br>ATGCGCTTCAAAGTCCACATGGAAGGCAGCGTTAATGGTCACGAGTTCGAAATTGAGG<br>GCGAAGGCGAAGGTCGTCCGTATGAGGGTACACAGACCGCTAAACTGAAAGTCACGA<br>AAGGTGGTCCACTGCCATTTGCTTGGGATATTCTGAGCCCACAGTTCATGTATGGCT<br>CCAAAGCCTATGTGAAACATCCGGCCGATATTCCGGACTATCTGAAACTGAGCTTCC<br>CTGAAGGGTTCAAATGGGAACGTGTGATGAACTTTGAGGATGGTGGTGTTGTGACAG<br>TGACACAGGATTCTAGCCTGCAAGACGGTGAGTTCATCTATAAAGTGAAACTGCGTG<br>GCACGAATTTTCCGAGTGATGGCCCGGTTATGCAGAAAAAAACGATGGGTTGGGAGG<br>CCTCTAGTGAGCGTATGTATCCAGAAGATGGCGCTCTGAAAGGCGAAATCAAACAGC<br>GTCTGAAACTGAAAGATGGTGGCCACTATGATGCCGAAGTGAAAACCACGTATAAAG<br>CCAAAAAACCTGTCCAACTGCCTGGTGCCTATAACGTTAACATCAAACTGGACATCA<br>CCTCACACAATGAGGACTATACGATCGTGGAGCAGTATGAGCGTGCTGAAGGACGTC<br>ATTCTACCGGTGGTATGGATGAGCTGTATAAATAATTCGAGCTCCGTCGACAAGC<br>TTTCTCAAAGCCCGCCGAAAGGCGGGCTTTTTTTT |

Table 3: Part sequences

| Name | Sequence [5'->3'] |
|---|---|
| Promoter J23119 | TTGACAGCTAGCTCAGTCCTAGGTATAATACTAGT |
| Promoter J23107 | tttacggctagctcagccctaggtattatgctagc |
| Promoter J23108 | ctgacagctagctcagtcctaggtataatgctagc |
| Promoter J23115 | tttatagctagctcagcccttggtacaatgctagc |
| Terminator t500 | CAAAGCCCGCCGAAAGGCGGGCTTTTTTTT |

| Name | Sequence [5'->3'] |
|---|---|
| Terminator TrrnB | GAAGCTTGGGCCCGAACAAAAACTCATCTCAGAAGAGGATCTGAATAGCGC<br>CGTCGACCATCATCATCATCATCATTGAGTTTAAACGGTCTCCAGCTTGGC<br>TGTTTTGGCGGATGAGAGAAGATTTTCAGCCTGATACAGATTAAATCAGAA<br>CCAGAAGCGGTCTGATAAAACAGAATTTGCCTGGCGGCAGTAGCGCGGTGG<br>TCCCACCTGACCCCATGCCGAACTCAGAAGTGAAACGCCGTAGCGCCGATG<br>GTAGTGTGGGGTCTCCCCATGCGAGAGTAGGGAACTGCCAGGCATCAAATA<br>AAACGAAAGGCTCAGTCGAAAGACTGGGCCTTTCGTTTTATCTGTTGTTTG<br>TCGGTGAACT |
| AmpR | CCAATGCTTAATCAGTGAGGCACCTATCTCAGCGATCTGTCTATTTCGTTC<br>ATCCATAGTTGCCTGACTCCCCGTCGTGTAGATAACTACGATACGGGAGGG<br>CTTACCATCTGGCCCCAGTGCTGCAATGATACCGCGAGACCCACGCTCACC<br>GGCTCCAGATTTATCAGCAATAAACCAGCCAGCCGGAAGGGCCGAGCGCAG<br>AAGTGGTCCTGCAACTTTATCCGCCTCCATCCAGTCTATTAATTGTTGCCG<br>GGAAGCTAGAGTAAGTAGTTCGCCAGTTAATAGTTTGCGCAACGTTGTTGC<br>CATTGCTACAGGCATCGTGGTGTCACGCTCGTCGTTTGGTATGGCTTCATT<br>CAGCTCCGGTTCCCAACGATCAAGGCGAGTTACATGATCCCCCATGTTGTG<br>CAAAAAAGCGGTTAGCTCCTTCGGTCCTCCGATCGTTGTCAGAAGTAAGTT<br>GGCCGCAGTGTTATCACTCATGGTTATGGCAGCACTGCATAATTCTCTTAC<br>TGTCATGCCATCCGTAAGATGCTTTTCTGTGACTGGTGAGTACTCAACCAA<br>GTCATTCTGAGAATAGTGTATGCGGCGACCGAGTTGCTCTTGCCCGGCGTC<br>AATACGGGATAATACCGCGCCACATAGCAGAACTTTAAAAGTGCTCAT |
| CmR | GGCACGTAAGAGGTTCCAACTTTCACCATAATGAAATAAGATCACTACCG<br>GGCGTATTTTTTGAGTTATCGAGATTTTCAGGAGCTAAGGAAGCTAAAAT<br>GGAGAAAAAAATCACTGGATATACCACCGTTGATATATCCCAATGGCATC<br>GTAAAGAACATTTTGAGGCATTTCAGTCAGTTGCTCAATGTACCTATAAC<br>CAGACCGTTCAGCTGGATATTACGGCCTTTTTAAAGACCGTAAAGAAAAA<br>TAAGCACAAGTTTTATCCGGCCTTTATTCACATTCTTGCCCGCCTGATGA<br>ATGCTCATCCGGAATTTCGTATGGCAATGAAAGACGGTGAGCTGGTGATA<br>TGGGATAGTGTTCACCCTTGTTACACCGTTTTCCATGAGCAAACTGAAAC<br>GTTTTCATCGCTCTGGAGTGAATACCACGACGATTTCCGGCAGTTTCTAC<br>ACATATATTCGCAAGATGTGGCGTGTTACGGTGAAAACCTGGCCTATTTC<br>CCTAAAGGGTTTATTGAGAATATGTTTTTCGTCTCAGCCAATCCCTGGGT<br>GAGTTTCACCAGTTTTGATTTAAACGTGGCCAATATGGACAACTTCTTCG<br>CCCCCGTTTTCACCATGGGCAAATATTATACGCAAGGCGACAAGGTGCTG<br>ATGCCGCTGGCGATTCAGGTTCATCATGCCGTTTGTGATGGCTTCCATGT<br>CGGCAGAATGCTTAATGAATTACAACAGTACTGCGATGAGTGGCAGGGCG<br>GGGCGTAATTTGATATCGAGCTCGCTTGGACTCCTGTTGATAGATCCAGT<br>AATGACCTCAGAACTCCATCTGGATTTGTTCAGAACGCTCGGTTGCCGCC<br>GGGCGTTTTTTATT |

| Name | Sequence [5'->3'] |
|---|---|
| ColE1 origin | GGCCGCGTTGCTGGCGTTTTTCCACAGGCTCCGCCCCCCTGACGAGCATC<br>ACAAAAATCGACGCTCAAGTCAGAGGTGGCGAAACCCGACAGGACTATAA<br>AGATACCAGGCGTTTCCCCCTGGAAGCTCCCTCGTGCGCTCTCCTGTTCC<br>GACCCTGCCGCTTACCGGATACCTGTCCGCCTTTCTCCCTTCGGGAAGCG<br>TGGCGCTTTCTCATAGCTCACGCTGTAGGTATCTCAGTTCGGTGTAGGTC<br>GTTCGCTCCAAGCTGGGCTGTGTGCACGAACCCCCCGTTCAGCCCGACCG<br>CTGCGCCTTATCCGGTAACTATCGTCTTGAGTCCAACCCGGTAAGACACG<br>ACTTATCGCCACTGGCAGCAGCCACTGGTAACAGGATTAGCAGAGCGAGG<br>TATGTAGGCGGTGCTACAGAGTTCTTGAAGTGGTGGCCTAACTACGGCTA<br>CACTAGAAGAACAGTATTTGGTATCTGCGCTCTGCTGAAGCCAGTTACCT<br>TCGGAAAAAGAGTTGGTAGCTCTTGATCCGGCAAACAAACCACCGCTGGT<br>AGCGGTGGTTTTTTTGTTTGCAAGCAGCAGATTACGCGCAGAAAAAAAGG<br>ATCTCAAGAAGATCCTTTGATCTTTTCTACGGGGTCTGACGCTCAGTGGA<br>ACGAAAACTCACGTTAAGGGATTTTGGTCATGA |
| p15A origin | GCGCTAGCGGAGTGTATACTGGCTTACTATGTTGGCACTGATGAGGGTGT<br>CAGTGAAGTGCTTCATGTGGCAGGAGAAAAAAGGCTGCACCGGTGCGTCA<br>GCAGAATATGTGATACAGGATATATTCCGCTTCCTCGCTCACTGACTCGC<br>TACGCTCGGTCGTTCGACTGCGGCGAGCGGAAATGGCTTACGAACGGGGC<br>GGAGATTTCCTGGAAGATGCCAGGAAGATACTTAACAGGGAAGTGAGAGG<br>GCCGCGGCAAAGCCGTTTTTCCATAGGCTCCGCCCCCCTGACAAGCATCA<br>CGAAATCTGACGCTCAAATCAGTGGTGGCGAAACCCGACAGGACTATAAA<br>GATACCAGGCGTTTCCCCCTGGCGGCTCCCTCGTGCGCTCTCCTGTTCCT<br>GCCTTTCGGTTTACCGGTGTCATTCCGCTGTTATGGCCGCGTTTGTCTCA<br>TTCCACGCCTGACACTCAGTTCCGGGTAGGCAGTTCGCTCCAAGCTGGAC<br>GTATGCACGAACCCCCCGTTCAGTCCGACCGCTGCGCCTTATCCGGTAAC<br>TATCGTCTTGAGTCCAACCCGGAAAGACATGCAAAAGCACCACTGGCAGC<br>AGCCACTGGTAATTGATTTAGAGGAGTTAGTCTTGAAGTCATGCGCCGGT<br>TAAGGCTAAACTGAAAGGACAAGTTTTGGTGACTGCGCTCCTCCAAGCCA<br>GTTACCTCGGTTCAAAGAGTTGGTAGCTCAGAGAACCTTCGAAAAACCGC<br>CCTGCAAGGCGGTTTTTTCGTTTTCAGAGCAAGAGATTACGCGCAGACCA<br>AAACGATCTCAAGAAGATCATCTTATTAATCAGATAAAATATTT |

| **Name** | **Sequence [5'->3']** |
|---|---|
| sfGFP | ATGAGCAAAGGAGAAGAACTTTTCACTGGAGTTGTCCCAATTCTTGTTGA<br>ATTAGATGGTGATGTTAATGGGCACAAATTTTCTGTCCGTGGAGAGGGTG<br>AAGGTGATGCTACAAACGGAAAACTCACCCTTAAATTTATTTGCACTACT<br>GGAAAACTACCTGTTCCGTGGCCAACACTTGTCACTACTCTGACCTATGG<br>TGTTCAATGCTTTTCCCGTTATCCGGATCACATGAAACGGCATGACTTTT<br>TCAAGAGTGCCATGCCCGAAGGTTATGTACAGGAACGCACTATATCTTTC<br>AAAGATGACGGGACCTACAAGACGCGTGCTGAAGTCAAGTTTGAAGGTGA<br>TACCCTTGTTAATGTATCGAGTTAAAGGGTATTGATTTTAAAGAAGATGG<br>AAACATTCTTGGACACAAACTCGAGTACAACTTTAACTCACACAATGTAT<br>ACATCACGGCAGACAAACAAAAGAATGGAATCAAAGCTAACTTCAAAATT<br>CGCCACAACGTTGAAGATGGTTCCGTTCAACTAGCAGACCATTATCAACA<br>AAATACTCCAATTGGCGATGGCCCTGTCCTTTTACCAGACAACCATTACC<br>TGTCGACACAATCTGTCCTTTCGAAAGATCCCAACGAAAAGCGTGACCAC<br>ATGGTCCTTCTTGAGTTTGTAACTGCTGCTGGGATTACACATGGCATGGA<br>TGAGCTCTACAAA |
| MGA4x | TCTAGATGGTGTTTTGGTTTGGTCAACGGATCCCGACTGGCGAGAGCCAG<br>GTAACGAATGGATCCGTTGACACCCAACAAAAAAAAACACGGATCCCGAC<br>TGGCGAGAGCCAGGTAACGAATGGATCCGTGTAAAAAAAAACCCAAAAAG<br>CCGGGATCCCGACTGGCGAGAGCCAGGTAACGAATGGATCCCGGCAACCA<br>ACCAACCAACCAAAAAGGATCCCGACTGGCGAGAGCCAGGTAACGAATGG<br>ATCCTTTTTCTAGCGCGGATCCGAATTCGAGCTCCGTCGACAAGCTT |
| Csy4 | ATGGACCACTACCTCGACATTCGCTTGCGACCGGACCCGGAATTTCCCCCG<br>GCGCAACTCATGAGCGTGCTCTTCGGCAAGCTCCACCAGGCCCTGGTGGCA<br>CAGGGCGGGGACAGGATCGGCGTGAGCTTCCCCGACCTCGACGAAAGCCGCT<br>CCCGGCTGGGCGAGCGCCTGCGCATTCATGCCTCGGCGGACGACCTTCGTGC<br>CCTGCTCGCCCGGCCCTGGCTGGAAGGGTTGCGGGACCATCTGCAATTCGGA<br>GAACCGGCAGTCGTGCCTCACCCCACACCGTACCGTCAGGTCAGTCGGGTTC<br>AGGCGAAAAGCAATCCGGAACGCCTGCGGCGGCGGCTCATGCGCCGGCACGA<br>TCTGAGTGAGGAGGAGGCTCGGAAACGCATTCCCGATACGGTCGCGAGAGCC<br>TTGGACCTGCCCTTCGTCACGCTACGCAGCCAGAGCACCGGACAGCACTTCC<br>GTCTCTTCATCCGCCACGGGCCGTTGCAGGTGACGGCAGAGGAAGGAGGATT<br>CACCTGTTACGGGTTGAGCAAAGGAGGTTTCGTTCCCTGGTTCTGA |

| **Name** | **Sequence [5'->3']** |
|---|---|
| dCas9 | GATAAGAAATACTCAATAGGCTTAGCTATCGGCACAAATAGCGTCGGATGGGC<br>GGTGATCACTGATGAATATAAGGTTCCGTCTAAAAAGTTCAAGGTTCTGGGAA<br>ATACAGACCGCCACAGTATCAAAAAAAATCTTATAGGGGCTCTTTTATTTGA<br>CAGTGGAGAGACAGCGGAAGCGACTCGTCTCAAACGGACAGCTCGTAGAAGGT<br>ATACACGTCGGAAGAATCGTATTTGTTATCTACAGGAGATTTTTTCAAATGAG<br>......................................................<br>CAATACGTGAACAAGCAGAAAATATTATTCATTTATTTACGTTGACGAATCTTGG<br>AGCTCCCGCTGCTTTTAAATATTTTGATACAACAATTGATCGTAAACGATATACG<br>TCTACAAAAGAAGTTTTAGATGCCACTCTTATCCATCAATCCATCACTGGTCTTTA<br>TGAAACACGCATTGATTTGAGTCAGCTAGGAGGTGAC |
| Target6 | CCAGTCATCAAGTCAGTCCAGTCAAAGTTTCCGTCGTTCAGCGGGGAATGT<br>ATACAGTTCATGTATATATTCCCCGCTTTTTTTTT |
| Target1 | CCATCTTACCTTTGCATCTCTATCGTTCTCAT<br>CTCATCCTGCGGGGAATGTATACAGTTCATGTATATATTCCCCGCTTTTTTTTT |
| Toehold2 | GGGAGTTTGATTACATTGTCGTTTAGTTTAGTGATACATAAACAGAGGAGA<br>TATCACATGACTAAACGAAACCTGGCGGCAGCGCAAAAG |
| Toehold3 | GGGATCTATTACTACTTACCATTGTCTTGCTCTATACAGAAACAGAGGAGA<br>TATAGAATGAGACAATGGAACCTGGCGGCAGCGCAAAAG |
| STAR AD1.A5 | TGAACTGTATACATTCCCCGCTGCTCCAACATTTATACAACTAATTAAAAC<br>AATTCACTGTAAAAACT |
| STAR6 | TGAACTGTATACATTCCCCGCTGAACGACGGAAACTTTGACTGGACTGACT<br>TGATGACTGG |
| STAR 1 | TGAACTGTATACATTCCCCGCAGGATGAGATGAGAA<br>CGATAGAGATGCAAAGGTAAGATGG |
| trigger2 | GGGACAGATCCACTGAGGCGTGGATCTGTGAACACTAAACTAAACGACAAT<br>GTAATCAAACTAAC |
| trigger3 | GGGTGATGGGACATTCCGATGTCCCATCAATAAGAGCAAGACAATGGTAAG<br>TAGTAATAGATAAG |
| SgRNA1 | tcctaggtataatactagttgtagtgacaagtgttggccaGT<br>TTTAGAGCTAGAAATAGCAAGTTAAAATAAGGCTAGTCCGTTATCAACTT<br>GAAAAAGTGGCACCGAGTCGGTGCTTTTTTTggatctcaaagcccgccgaa |
| SgRNA2 | gtagtgacaagtgttggccaGTTTTAGAGCTAGAAATAGCAA GTTAAAATAAGGCTAGTCCGTTAT-<br>CAACTTGAAAAAGTGGCACCGAGTCGGTGCTTTTTTT |
| SgRNA3 | caaactcaagaaggaccatgGTTTTAGAGCTAGAAATAGCAAGTTAAAATA<br>AGGCTAGTCCGTTATCAACTTGAAAAAGTGGCACCGAGTCGGTGCTTTTTTT |
| SgRNA4 | GGGACAGATCCACTGAGGCGTGGATCTGTGAACACTAAACTAAACGACAAT<br>GTAATCAAACTAAC |
| Csy4 hairpin | GTTCACTGCCGTATAGGCAGCTAAGAAA |

| **Name** | **Sequence [5'->3']** |
|---|---|
| Ion UTR | TTAAACTAAGAGAGAGCTCT |
| RplY+SD UTR | TTAAACTAAGGAGGAGCTCT |
| T7 UTR | gctagcaataattttgtttaacttta |
| ybaB+SD UTR | TTAAACTAACACACAGCTCT |
| ybaB-AG UTR | CGTGATTCACACAGAAACCT |
| Irp UTR | AATACAGAGAGACAATAATT |
